# Oxytocin receptor controls promiscuity and development in prairie voles

**DOI:** 10.1101/2024.09.25.613753

**Authors:** Ruchira Sharma, Kristen M. Berendzen, Amanda Everitt, Belinda Wang, Gina Williams, Shuyu Wang, Kara Quine, Rose D. Larios, Kimberly L. P. Long, Nerissa Hoglen, Bibi Alika Sulaman, Marie C. Heath, Michael Sherman, Robert Klinkel, Angela Cai, Denis Galo, Lizandro Chan Caamal, Nastacia L. Goodwin, Annaliese Beery, Karen L. Bales, Katherine S. Pollard, Arthur Jeremy Willsey, Devanand S. Manoli

**Affiliations:** Department of Psychiatry and Behavioral Sciences, University of California, San Francisco; San Francisco, CA 95158, USA; Center for Integrative Neuroscience, University of California, San Francisco; San Francisco, CA 95158, USA; Weill Institute for Neurosciences, University of California, San Francisco; San Francisco, CA 95158, USA; Kavli Institute for Fundamental Neuroscience, University of California, San Francisco; San Francisco, CA 95158, USA; Neurosciences Graduate Program, University of California, San Francisco; San Francisco, CA 95158, USA; Department of Integrative Biology, UC Berkeley, Berkeley, CA, United States; Department of Neuroscience, UC Berkeley, Berkeley, CA, United States; Department of Psychology, University of California, Davis; Davis, CA 95616, USA; Department of Neurobiology, Physiology, and Behavior, University of California, Davis; Davis, CA 95616, USA; Gladstone Institutes, San Francisco, CA, USA; Department of Epidemiology & Biostatistics, University of California, San Francisco, San Francisco, CA, USA; Chan Zuckerberg Biohub, San Francisco, CA, USA; Calico Labs, San Francisco, CA 94080, USA; Neurona Therapeutics, South San Francisco, CA 94080, USA; Department of Neuroscience, University of Michigan, Ann Arbor, MI 48109, USA; Department of Neurobiology and Behavior, University of California, Irvine; Irvine, CA 92697, USA; Branch Out School, Los Angeles, CA 90026, USA; University of California, Los Angeles; Los Angeles, CA 90095, USA; Department of Psychology, University of Washington; Seattle, WA 98195, USA; Department of Rehabilitation and Regenerative Medicine, Columbia University Irving Medical Center; New York, NY 10032, USA

**Keywords:** Attachment behaviors, prairie vole, pair-bonding, oxytocin receptor, partner preference, monogamy, promiscuity, oxytocin signaling, paraventricular nucleus of the hypothalamus, nucleus accumbens

## Abstract

Oxytocin receptor (Oxtr) signaling influences complex social behaviors in diverse species, including social monogamy in prairie voles. How Oxtr regulates specific components of social attachment behaviors and the neural mechanisms mediating them remains unknown. Here, we examine prairie voles lacking Oxtr and demonstrate that pair bonding comprises distinct behavioral modules: the preference for a bonded partner, and the rejection of novel potential mates. Our longitudinal study of social attachment shows that Oxtr sex-specifically influences early interactions between novel partners facilitating the formation of partner preference. Additionally, Oxtr suppresses promiscuity towards novel potential mates following pair bonding, contributing to rejection. Oxtr function regulates coordinated patterns of gene expression in regions implicated in attachment behaviors and regulates the expression of oxytocin in the paraventricular nucleus of the hypothalamus, a principal source of oxytocin. Thus, Oxtr controls genetically separable components of pair bonding behaviors and coordinates development of the neural substrates of attachment.

## Introduction

Social attachments, such as the bonds between parents and children, family and kin, friends, and the long-term relationships between mates and romantic partners form the basis of interpersonal and group dynamics within societies^1–7^. Across mammals and other taxa, various selective pressures have given rise to diverse patterns of complex social organization involving long-term relationships between members of a species, one of the most robust and intriguing of which is the enduring bond between mates^2,4,8–13^. In mammals, ∼4% of species demonstrate social monogamy, forming persistent pair bonds between mating partners^9,14–16^. Comparative studies of closely related species that display distinct patterns of social behaviors, including social monogamy, reveal correlated species-specific differences in the nonapeptide hormones oxytocin (Oxt) and arginine vasopressin (Avp), or their homologs, suggesting that these hormones serve as critical modulators of affiliative behaviors and pair bonding^9–11,16–20^. Despite the central role of social attachment in human behavior and a rich history of investigation into the cognitive processes underlying social cognition and affiliative behaviors, it has been difficult to probe the neural mechanisms underlying these fascinating behaviors at the cellular, molecular, and genetic level, as most species that serve as model organisms for such studies in the laboratory do not display long-term social attachment as adults ^6,21–27^.

Prairie voles (*Microtus ochrogaster*) display long-term social attachment such that mating partners show an enduring pair bond and social monogamy^13,28–31^. Like many complex innate behaviors, pair bonding consists of distinct modules: the development of increased prosocial behaviors with partners over strangers (partner preference), and increased agonistic behaviors towards novel opposite sex potential mates (stranger rejection)^13,30,32–37^. As in humans and other species that show affiliative social behaviors, disruption of pair bonds results in increased anxiety-like behaviors and markers of stress, supporting integrated neural and physiologic mechanisms that facilitate the preservation of such attachments^38–40^. Pioneering work in prairie voles identified Oxt and Avp as critical mediators of pair bonding^41,42^. Interspecific variations in the patterns of expression of the oxytocin receptor (Oxtr) and the vasopressin 1a receptor (V1ar) correlate with the potential for pair bonding within closely related vole species^43–49^.

Across phyla, manipulations of oxytocin or its species-specific homolog alters social behaviors. In prairie voles, pharmacologic inhibition of Oxt and Avp signaling via their respective cognate receptors disrupts pair bonding, while exogenous administration of these hormones promotes pair bonding without mating^28,50,51^. Transient blockade of Oxtr only during the first few hours of cohabitation delays the formation of a partner preference^28,29^ and artificial overexpression of Oxtr accelerates the formation of a pair bond^52^. Exogenous oxytocin administration leads to to increased prosocial behaviors towards conspecifics as well as humans in domestic dogs, initiation of huddling towards a potential partner in marmosets (*Callithrix penicillata*), and food sharing with conspecifics in pinyon jays (*Gymnorhinus cyanocephalus*, a highly social corvid species)^53–56^. Conversely, OT receptor antagonists administered during early pair-bond formation to monogamous cichlid fish (*Amatitlania nigrofasciata*), socially monogamous zebra finch (*Taeniopygia guttata*) and marmosets (*Callithrix penicillata*) lead to reduced affiliative behaviors towards potential mates^55,57,58^. Taken together, these studies underscore the broad conservation of this neuropeptide in the modulation of social behaviors, as well an early temporal window during which oxytocin-dependent behaviors mediate partner preference.

Given the central role of Oxt signaling in affiliative behaviors, we tested the genetic requirement for signaling via Oxtr for pair bonding and social behaviors in prairie voles. We generated prairie voles lacking Oxtr and unexpectedly observed that these animals continue to display a strong preference for partners following mating and cohabitation^59^. To gain deeper insight into the modulation of attachment and social behaviors by Oxtr function, we developed a series of behavioral assays to probe distinct aspects of pair bonding and associated social behaviors. To interrogate the molecular mechanisms mediating social attachment, we profiled patterns of gene expression in the nucleus accumbens (NAc), a key site of species differences associated with social monogamy that is dramatically enriched for Oxt binding in prairie voles^45,60,61^. We find that Oxtr controls the timing of pair bonding in prairie voles and promiscuity independent of partner preference formation. Oxtr sex-specifically influences the behavior of potential mates, increasing agonistic displays in naive wildtype (WT) females, evidenced by decreased aggression and increased prosocial interactions shown by potential female WT mates in the presence of a male lacking Oxtr. Oxtr function regulates patterns of gene expression in the NAc, as well as other regions of the vole brain implicated in social and attachment behaviors, in distinct ways in each sex. Furthermore, we see changes in expression of genes regulating neurodevelopmental processes and disease, supporting a role for Oxtr function early in life^62–64^. Finally, consistent with potential trophic effects of Oxtr function during development, we find that loss of Oxtr alters patterns of Oxt and Avp expression in the paraventricular nucleus, a principal source of these neuropeptides^63,65^. Our studies reveal genetically separable components of pair bonding and suggest that Oxtr signaling sex-specifically participates in the development of the circuits underlying attachment behaviors.

## Results

### Oxtr function facilitates the formation of partner preference

We previously found that partner preference formation, following 1 week of cohabitation, can occur in the absence of Oxtr function^59^. However, acute modulation of Oxtr signaling can alter patterns of social behaviors between individuals^29,50,66^. These observations suggest that changes in Oxtr function may influence early social interactions that contribute to the trajectory of pair bond formation and/or specific aspects of attachment behaviors following the formation of partner preference. We first wished to determine whether Oxtr function facilitated pair bond formation by examining the display of partner preference following shorter periods of cohabitation. Consistent with previous studies showing that the cohabitation time required for the display of partner preference differs between males and females, we find that WT females demonstrate partner preference after only 6 hours of cohousing with a WT male partner, while WT males require 5 days following introduction to a potential WT mate (Fig. S1A-C)^67^. We therefore cohoused WT or Oxtr^1-/-^ females with WT males for 6 hours, and WT or Oxtr^1-/-^ males with WT females for 5 days following introduction (short cohabitation), or in these combinations for 1 or 7 days (long cohabitation) for females or males, respectively (Fig. 1A, Fig. S1A).

**Figure 1:**
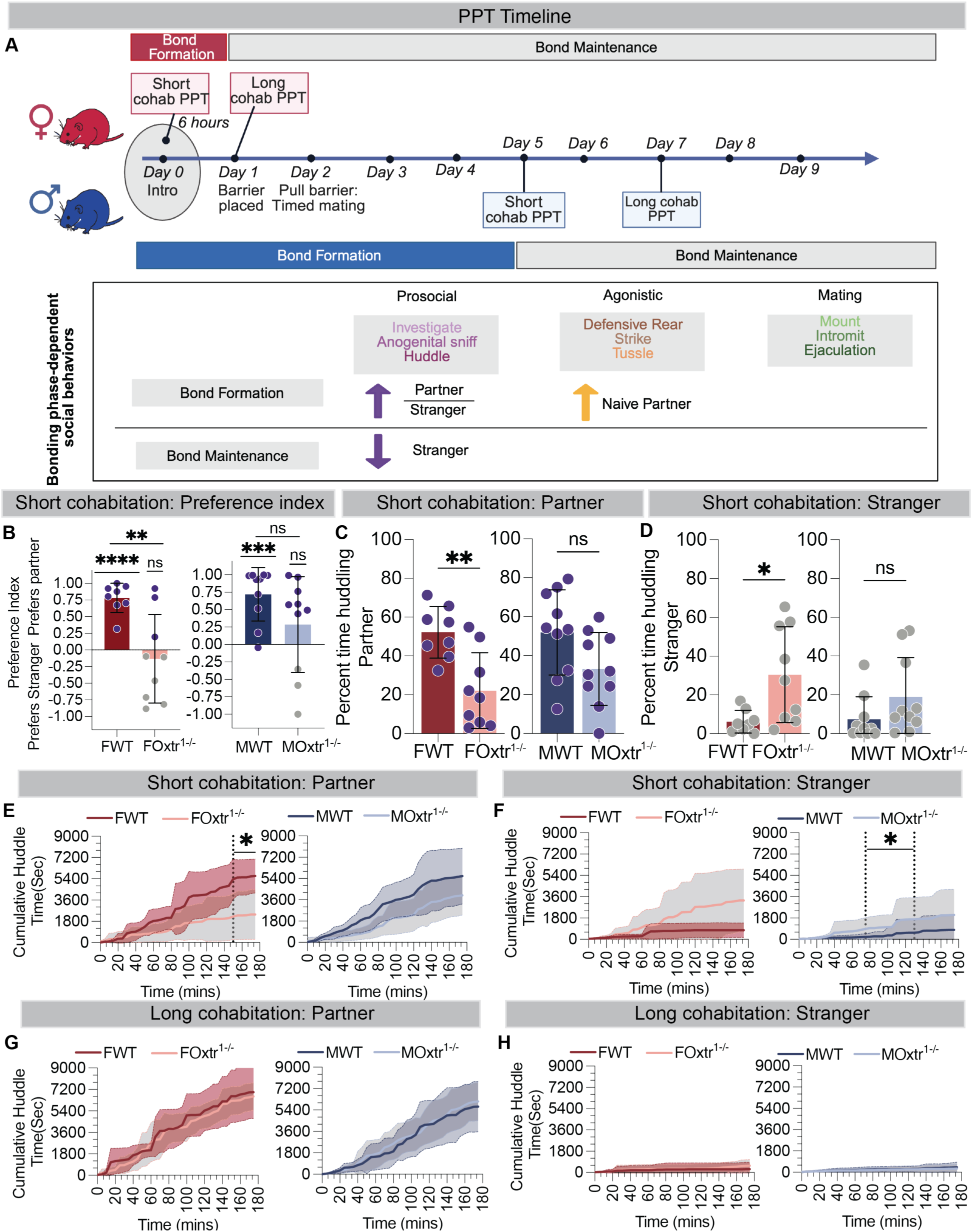
Oxtr reduces the amount of time naive animals take to show partner preference. A. Schematic of the partner preference tests (PPTs) and the day on which they were conducted to study pair bonding behaviors. The prosocial, agonistic and mating behaviors scored are shown below. During the bond formation phase, the ratio of partner-directed vs stranger-directed affiliative behaviors increases in wild types and following the bond formation in the bond maintenance phase, wild types show decreased prosocial behaviors and increased agonistic behaviors towards strangers. B. A preference for partner is induced in the WTs but not Oxtr^1-/-^ animals following short cohabitation. C. WT females spend more time huddling with their partners. D. Oxtr^1-/-^ females spend more time huddling with strangers. E. WT females show a significant increase in partner huddling time towards the end of a 3-hour assay. F. Oxtr^1-/-^ males show a significant increase in stranger huddling time in the middle of a 3-hour assay. G. No difference in partner huddling time between WT and Oxtr^1-/-^ animals following long cohabitation. H. No difference in stranger huddling time between WT and Oxtr^1-/-^ animals following long cohabitation. See Supplementary Figure 1 and Supplementary Table 1. Mean +/− SD, n =>8, *p =< 0.05, FWT = female WT, FOxtr^1-/-^ = female Oxtr^1-/-^, MWT = male WT, MOxtr^1-/-^ = male Oxtr^1-/-^.

Both males and females lacking Oxtr show an absence of partner preference following short cohabitation with WT partners (Fig. 1B). Oxtr^1-/-^ females, but not males, showed a decrease in the fraction of huddling time with their partner and an increase in the fraction of huddling time with a stranger compared to WT controls (Fig. 1C-D). Oxtr^1-/-^ males present an absence of partner preference (Fig. 1B), resulting from a trend towards decreased time with their cohoused partner and increased time with strangers (Fig. 1C-D). Consistent with our previous studies, we find no difference between WT and Oxtr^1-/-^females when the pairs are cohoused for the longer duration (Fig. S1B-F).

To better understand the social interaction dynamics underlying differences in huddling behavior between WT animals and their siblings lacking Oxtr after short-term cohabitation, we quantified the cumulative time subjects spent huddling with either their cohoused partners or strangers across the assay.Oxtr^1-/-^ females show a significant decrease in the amount of time spent huddling with partners in the last 30 mins of the assay (150-180mins) (Fig. 1E). We also observe a significant increase in huddling time of Oxtr^1-/-^ males with strangers during the middle periods (75 – 130mins) (Fig. 1F), suggesting that they engage in more early prosocial behavior with strangers than WT males. Consistent with our previous work, we find no difference between cumulative huddling times with either partner or stranger between WTs and Oxtr^1-/-^ animals of both sexes after long cohabitation periods (Fig. 1G-H)^45^. These observations demonstrate that Oxtr function changes the temporal dynamics of social interactions with both familiar and unfamiliar animal in a sex-specific manner. Taken together, our findings suggest that Oxtr function acts early in the process of pair bonding to facilitate the formation of the partner preference.

### Oxtr suppresses promiscuous behaviors in a state dependent manner in females

Oxtr influences the dynamics of interactions with partners and strangers differently between the sexes. We therefore wished to examine the detailed patterns of these interactions separately in females and males (Fig. 2A). We examined the distribution of huddles by WT vs Oxtr^1-/-^ females following short cohabitation as a measure of prosocial behavior displayed by these animals. We find that the top quartile of huddle durations (long huddles) from each subject animal make up an average of ∼90% of the total huddle duration, functionally separating these interactions from incidental side-by-side contact that may not constitute affiliative huddling (Fig. S2A Left). We classified long huddles as partner huddles or stranger huddles and discovered that, consistent with the overall time spent, the median huddle duration carried out by Oxtr^1-/-^ females is significantly shorter with partners and significantly longer with strangers, compared to WT females (Fig. 2B). The longest huddle displayed with partner is significantly longer for WT females compared to Oxtr^1-/-^ females though we find no difference in the length of the overall longest huddle. This is due to some Oxtr^1-/-^ females conducting their longest huddle with the stranger instead (Fig. S2B). Oxtr^1-/-^ females additionally executed a larger number of huddles only with the stranger, but not the partner, compared to WT controls (Fig. 2C). Taken together, these results suggest that Oxtr function in females both facilitates prosocial behavior with partners and suppresses such behavior with strangers during the early stages of pair bond formation.

**Figure 2:**
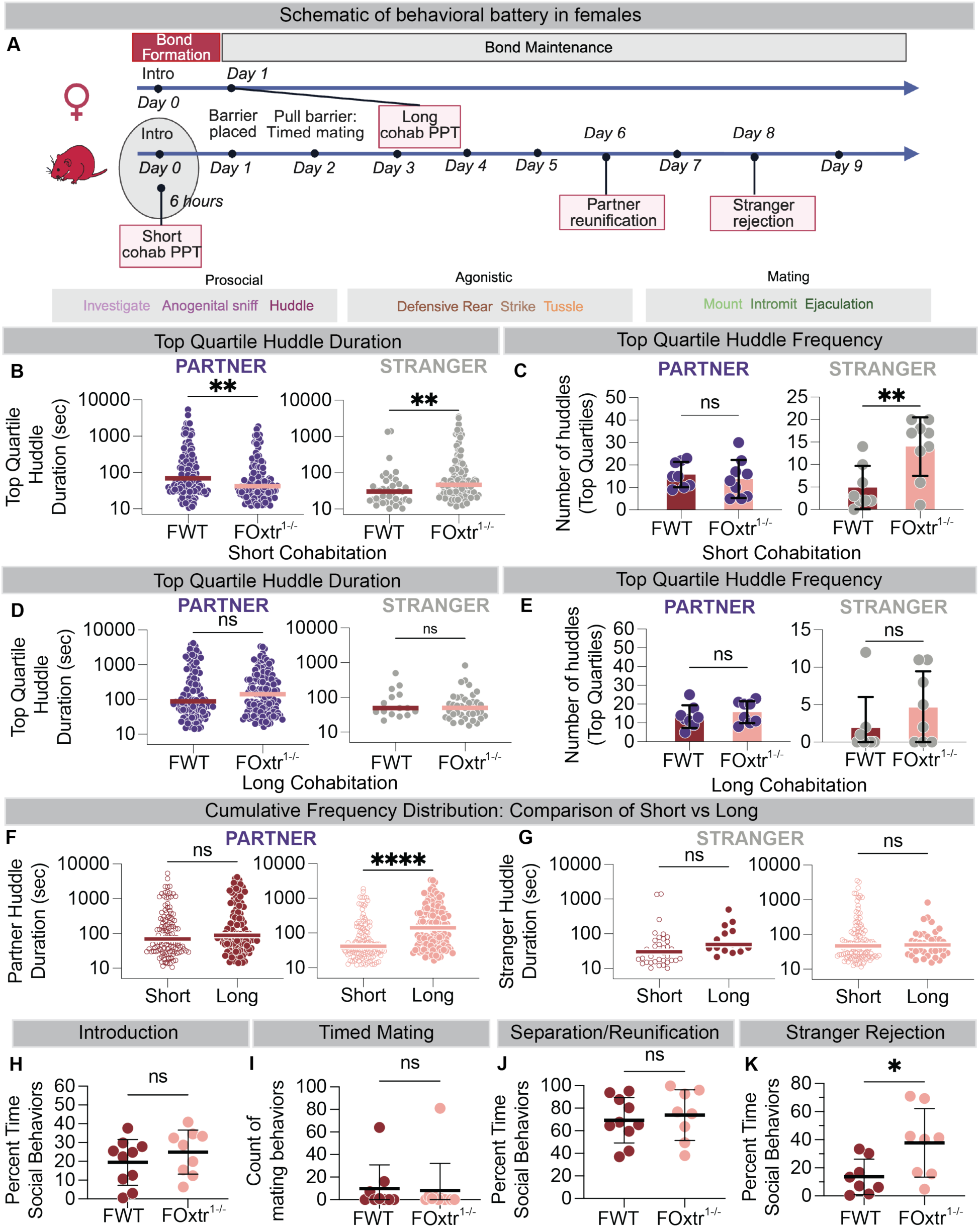
Oxtr suppresses promiscuous behaviors in a state dependent manner in females. A. Timeline of the behavioral battery carried out on females. Bottom row shows the behaviors scored. B. Oxtr^1-/-^ female long huddle duration is shorter with partners and longer with strangers after a short cohabitation. C. Oxtr^1-/-^ female long huddle frequency is greater with strangers, but not partners after a short cohabitation. D. No difference in median long huddle durations between WT and Oxtr^1-/-^ females post long cohabitation. E. No difference in long huddle frequencies between WT and Oxtr^1-/-^ females post long cohabitation. F. Increasing cohabitation time increases the median long huddle duration an Oxtr^1-/-^female executes with the partner. G. There is no significant change in the median long huddle duration conducted by females with strangers when cohabitation time is increased. H. Naive Oxtr^1-/-^ and WT females are equally prosocial towards a naive WT male. I. There is no difference in the counts of mating behaviors between Oxtr^1-/-^ and WT females. J. Oxtr^1-/-^ and WT females are equally prosocial towards their partners after a brief separation. K. Oxtr^1-/-^ females are significantly more prosocial towards a naive WT stranger male compared to WT females. See Supplementary Figure 2 and Supplementary Table 1. Mean ± SD, n =>8, *p < 0.05, FWT = female WT, FOxtr^1-/-^ = female Oxtr^1-/-^, MWT = male WT, MOxtr^1-/-^ = male Oxtr^1-/-^.

We next wished to determine whether the loss of Oxtr function influences patterns of prosocial behaviors after the display of partner preference. We therefore examined patterns of huddling, in particular the duration and frequency of these interactions, following long cohabitation, a time point at which both WT and Oxtr^1-/-^ animals display robust preference for partners over strangers^59^ (Fig. S1D). As with short cohabitation, the top quartile of huddles (by duration) for each subject animal following long cohabitation, described an average of 90% of total huddle time (Fig. S2A right). Following long cohabitation, preference and huddling behavior in Oxtr^1-/-^ females is indistinguishable from their WT siblings (Fig. 2D-E). Comparing changes in the frequency of top quartile huddles displayed by Oxtr^1-/-^ females after short vs long cohabitation, we find that both WT and Oxtr^1-/-^ females show decreased huddle frequency with stranger following long cohabitation (Fig. S2C-D). On comparing durations of long huddles, we find that Oxtr^1-/-^females significantly increase their median huddle duration with partner post long cohousing (Fig. 2F-G). Thus, loss of Oxtr function in females reduces the duration of early prosocial interactions (short cohabitation) with partners and strangers while Oxtr-independent mechanisms eventually promote WT patterns of affiliative huddling and partner preference following longer periods of cohabitation.

Our results suggest that Oxtr function influences specific aspects of early pair bonding and attachment behaviors. We next wished to dissociate the influence of Oxtr on the formation and display of partner preference from the display of agonistic rejection behaviors towards novel strangers following cohabitation. We therefore developed a battery of assays to examine initial social interactions between mating partners at early and later stages of pair bonding, as well as stranger rejection following cohabitation (Fig.2A). We examined the behavior of subject females from each genotype with WT males when naive animals were first introduced to each other (Introduction), mating behaviors following estrus induction (Timed Mating), prosocial interactions following reunification with their bonded mate (Partner Reunification), and agonistic rejection of novel, opposite sex strangers (Stranger Rejection) (Fig. 2A see Methods). We find no differences in social behaviors during Introduction, Timed Mating and Partner Reunification (Fig. 2H-J). In contrast, during Stranger Rejection, Oxtr^1-/-^ females show more social behaviors towards novel naive males than WT females do in the absence of their partner (Fig. 2K). Taken together, these studies demonstrate that Oxtr function regulates specific components of pair bonding in females in a state-dependent manner, suppressing promiscuous social interactions with stranger males after cohabitation sufficient for the display of partner preference.

### Oxtr function influences different prosocial behaviors across the stages of pair bonding in males

Given the Oxtr-dependent difference in huddling behavior between males and females after short cohabitation (Fig. 1C-D), we wished to assess sex differences in the patterns of behaviors displayed by Oxtr^1-/-^ animals and to further investigate Oxtr function in males (Fig. 3A). Examining long huddles (as above, Fig. 2), we find that such huddles by Oxtr^1-/-^ males with their partners following short cohabitation are significantly shorter when compared to WT siblings, contributing to diminished partner preference early in pair bonding (Fig. 3B Left). Interestingly, while Oxtr^1-/-^ males show no difference in the total amount of time spent huddling with strangers when compared to WT siblings (Fig. 1D), they display shorter and more frequent huddles towards them at this time (Fig. 3B, 3C Right). In males, Oxtr thus influences the pattern as well as amount of early affiliative behavior to distinct behavioral stimuli in different ways.

**Figure 3:**
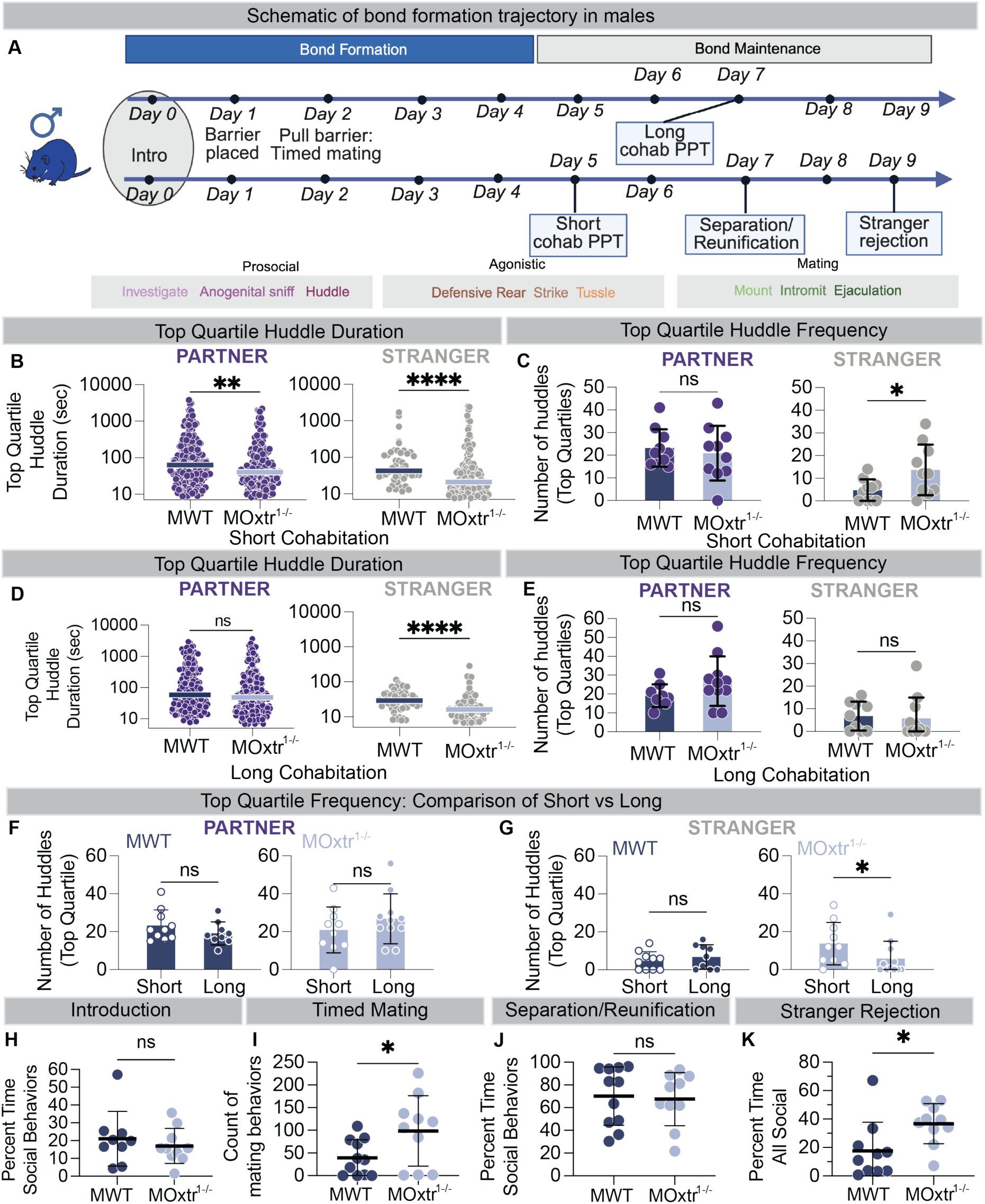
Oxtr function influences different prosocial behaviors across pair bonding in males. A. Timeline of the behavioral battery carried out on males. Bottom row shows the behaviors scored. B. Oxtr^1-/-^ male long huddle duration is shorter with partners and strangers after a short cohabitation. C. Oxtr^1-/-^ male long huddle frequency is greater with strangers, but not partners after a short cohabitation. D. The median long huddle executed by Oxtr^1-/-^ males with the stranger after a long cohabitation, is shorter than their WT siblings. E. No difference in the long huddle frequencies between WT and Oxtr^1-/-^ males post long cohabitation. F. There is no change in the frequency of long huddles with partner between Oxtr^1-/-^and WT males. G. Increasing cohabitation time decreases the frequency of long huddles with strangers in Oxtr^1-/-^ males. H. Naive Oxtr^1-/-^ and WT males are equally prosocial towards naive WT females. I. Oxtr^1-/-^ males execute higher counts of mating behaviors compared to WT. J. Prosocial behaviors following a brief separation with their partner are not different between Oxtr^1-/-^ and WT males. K. Oxtr^1-/-^ males are significantly more prosocial towards a naive WT stranger female compared to WT males. See Supplementary Figure 3 and Supplementary Table 1. Mean +/− SD, n =>8, *p =< 0.05, FWT = female WT, FOxtr^1-/-^ = female Oxtr^1-/-^, MWT = male WT, MOxtr^1-/-^ = male Oxtr^1-/-^

Oxtr^1-/-^ males show no difference in time spent with either partners or strangers after long cohabitation (Fig. S1E-F). However, these males show a persistent reduction in the duration of their long huddles with strangers at this time point (Fig. 3D). Following long cohabitation, WT and Oxtr^1-/-^ males show no difference in the duration of long huddles with partners and a reduction in the duration of long huddles with strangers (Fig. S3A-B). We find no difference in the frequency of long huddles between Oxtr^1-/-^ and WT males after long cohabitation (Fig. 3E). Consistently, comparing Oxtr^1-/-^ males’ behavior following short vs long cohabitation reveals that the frequency with which Oxtr^1-/-^ males huddle with strangers decreases following long cohabitation (Fig. 3F-G). Thus, specifically in males, loss of Oxtr function does not change the overall time spent huddling with strangers after short or long cohabitation but does cause a continuing disruption in the patterns of affiliative behavior that comprises huddling with strangers.

Our observations suggest that the loss of Oxtr disrupts certain characteristics of prosocial behaviors in males regardless of the time spent bonding with their partner. We next wished to dissect whether such loss of function resulted in atypical affiliative or agonistic behaviors across stages of bonding. We examined the behavior of Oxtr^1-/-^ males and their WT siblings in the same series of paradigms described previously to characterize the formation and maintenance of their pair bonds (Fig. 3A). Oxtr^1-/-^ males spend similar amounts of time as WT males engaging in prosocial behaviors with partners regardless of bonding state (Fig. 3H-J, S3E). Similar to our observations in Oxtr^1-/-^ females, Oxtr^1-/-^males spend more time engaging in prosocial behaviors with strangers in the absence of their partners (Fig. 3K).

In contrast to time spent conducting mating behaviors with WT female partners (Fig. S3E), Oxtr^1-/-^ males demonstrate increased frequency of mating attempts when these females are in estrus (Fig. 3I). Examining all instances of mounting, we find that Oxtr^1-/-^ males carry out shorter mounting bouts more frequently than WT males (Fig. S3F). To determine whether the shorter bouts of mounting in Oxtr^1-/-^ males influence the receptivity of their partners, we compared the mount-to-intromission ratio for each animal^68^ and find that females are equally receptive to Oxtr^1-/-^ males as their WT siblings (Fig. S3G). Thus, consistent with our previous observations, Oxtr influences patterns, but not the amount or success, of male mating behaviors, suggesting that such changes in males’ behaviors are equally able to elicit receptivity in females.

Oxtr function influences distinct components of males’ attachment-related social behaviors towards unfamiliar females at different stages of pair-bonding, lengthening mating bout durations when naïve and suppressing promiscuous prosocial behaviors upon bonding. Taken together our results suggest that Oxtr influences patterns of social behaviors with future partners during the early phases of pair-bonding differently in females and males, but upon bond formation, functions in both sexes to suppress promiscuous affiliative behaviors towards strangers (Fig. S3H).

### Oxtr sex-specifically influences social interactions between potential mates

Oxtr function influences patterns of social interactions between mates early during the formation of a pair-bond (Short Cohabitation in both sexes, Introduction and Timed Mating in males). We therefore investigated the effect of Oxtr loss on the choice made by a potential mate. We developed a variation of the PPT three-chamber assay to present a naive WT “chooser” with a choice between a naive opposite sex WT or naive opposite sex Oxtr^1-/-^ animal (Fig. 4A, Fig. S4A, see Methods). Using the “naïve-choice” paradigm we find that, over the course of 6 hours, WT choosers spend significantly more time with one of the two stimulus animals (winner) (Fig. 4B) and that WT and Oxtr^1-/-^ animals are chosen as the winner with equal frequency(Fig. 4C).

**Figure 4:**
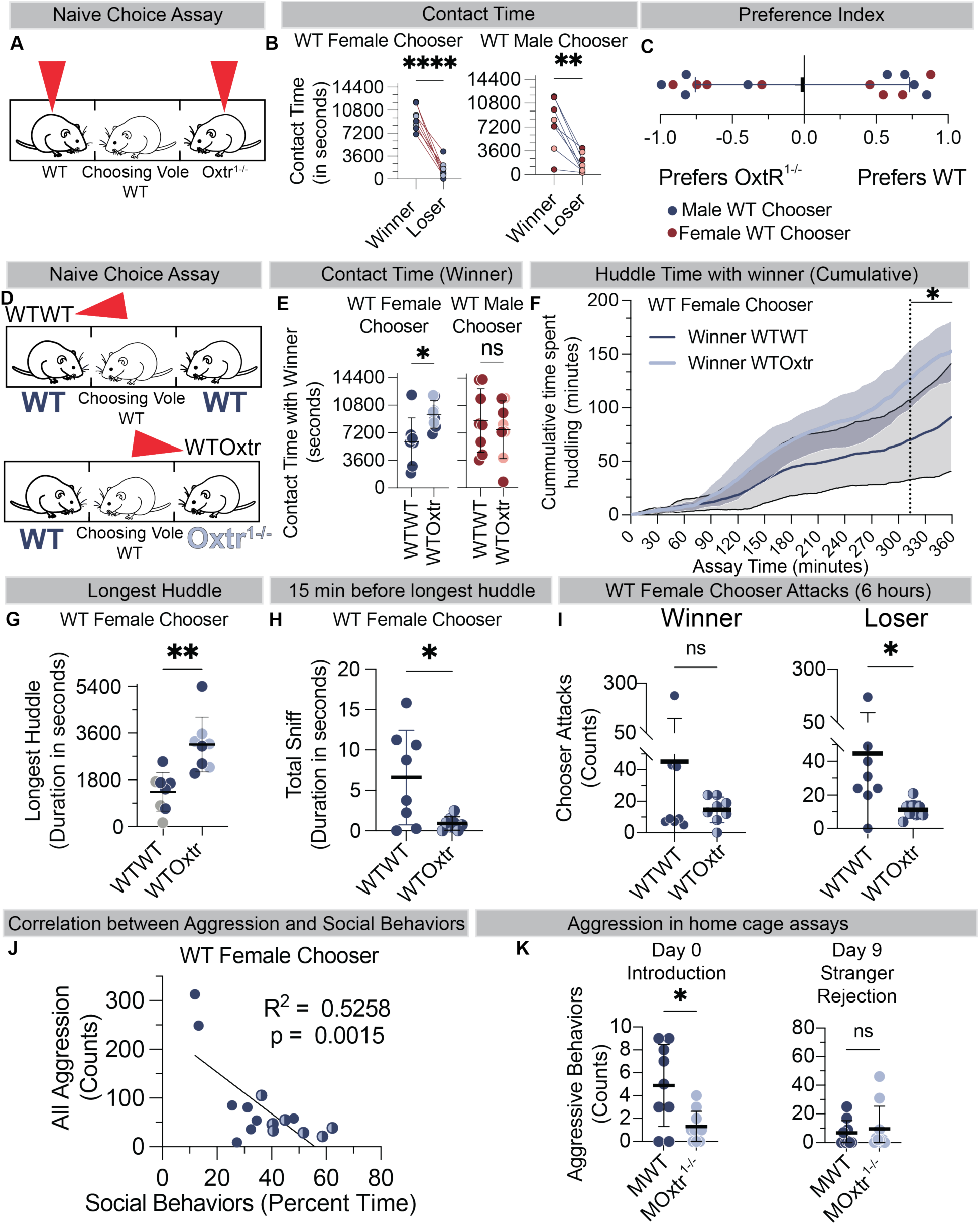
Oxtr sex-specifically influences social interactions between potential mates. A. Schematic of naive social choice with a naive WT animal choosing between an opposite sex WT and opposite sex Oxtr^1-/-^ animal (red triangles). B. WT choosers of both sexes spend significantly more time with one stimulus animal (winner) over the other (loser). The color of each dot denotes the genotype of the winner and loser. C. WT and Oxtr^1-/-^ animals are chosen as winners with equal frequency by WT choosers of both sexes. D. Schematic of comparison of contact with winner from WTOxtr condition to that with winner from WTWT condition. E. Female WT choosers spends more time in contact with a winner from a WTOxtr condition than the WTWT. The color of the dots represents the genotype of the winner. F. Female WT choosers spend significantly more time huddling with the winner from the WTOxtr condition in the last 47 mins of the assay. Solid lines depict the average cumulative huddle time, the solid black bar denotes the time at which the lines significantly diverge. G. Longest huddle from WTOxtr lanes is significantly higher compared to those from WTWT lanes. The color of each dot denotes the genotype/position of the animal with whom the longest huddle took place. H. Female WT choosers from WTWT lanes spend more time sniffing both stimulus animals 15 minutes before the longest huddle compared to female choosers from WTOxtr lanes I. Female WT choosers attack the loser in WTWT lanes more frequently than choosers from WTOxtr lanes (right). There is no difference in the number of attacks on the winners by female choosers from WTWT lanes compared to those from WTOxtr lanes (left). J. Total prosocial behavior time shows significant negative correlation with the total counts of aggressive behaviors displayed by female WT choosers. K. Naive male WTs attack a potential female WT mate more frequently than male Oxtr^1-/-^ animals (left). There is no difference in the number of attacks against an unfamiliar naive WT female stranger by bonded WT and Oxtr^1-/-^ males (right). See Supplementary Figure 4 and Supplementary Table 1. Mean +/− SD, n = 8, *p =< 0.05, FWT = female WT, FOxtr^1-/-^ = female Oxtr^1-/-^, MWT = male WT, MOxtr^1-/-^ = male Oxtr^1-/-^, WTWT = Lane with 2 naive WT animals, WTOxtr = Lane with a WT and a Oxtr^1-/-^ animal.

To examine whether Oxtr influences the dynamics of the choice made by a potential WT mate, we compared the time spent with the winner from a lane where the chooser had a choice between a naïve WT or Oxtr^1-/-^ animal (WTOxtr lanes) versus that of a chooser choosing between 2 naive WT animals (WTWT lanes) of the opposite sex (Fig. 4D). While naïve WT choosers show no difference in overall social choice between naive WT and Oxtr^1-/-^ opposite sex animals (Fig. 4C), WT female choosers show distinct behavioral dynamics in the presence of an Oxtr^1-/-^ male (WTOxtr lane). In the presence of an Oxtr mutant (WTOxtr lane), WT females spend significantly more time in contact with the winner, regardless of genotype, when compared to contact with the winner between two WT males (WTWT lane) (Fig. 4E). In contrast, we observe no difference in contact time with a winner when a naive WT male is choosing in WTOxtr versus WTWT lane (Fig. 4E).

We therefore manually examined the specific behaviors shown by female WT choosers during their interactions with both the winner and loser in WTWT and WTOxtr lanes. Preference indices using these metrics were highly concordant with automated scoring (Fig. S4A). The time spent by the female chooser displaying prosocial behaviors towards the winner and the time spent huddling with the winner in WTOxtr lanes was significantly greater than WTWT lanes (Fig. S4B) (Fig. 4E). The divergence between the amount of cumulative time spent huddling with the winner from WTOxtr versus WTWT lanes occurs after 5 hours have elapsed (313-360 mins, Fig. 4F), suggesting that complex interactions in the presence of a male lacking Oxtr influence how females “choose” a male partner regardless of whether they end up choosing a WT or Oxtr^1-/-^ winner.

We evaluated the duration of the longest huddles conducted by all WT female choosers, and find that the longest huddle is always conducted with the winner. The longest huddles conducted in the WTOxtr lanes, regardless of the genotype of the male winner, are greater than those from WTWT lanes (Fig. 4G). There is no difference in the latency to start the longest huddle between WTOxtr and WTWT lanes (Fig. S4C). Thus, the presence of the Oxtr^1-/-^ male in the lane leads a WT chooser female to execute the longest huddle for a greater amount of time compared to the longest huddle by the WT female chooser from a WTWT lane (Fig. 4G). These observations suggests either that naïve Oxtr^1-/-^ males elicit longer longest-huddle bouts from WT females, even when this huddle occurs with a WT male winner, or that the behavior of naïve WT males reduces the duration of WT females’ longest huddle - a reduction that is absent in the presence of a male lacking Oxtr.

To determine if specific patterns of behavior preceded the longest huddle with a winner, we aligned the onset of the longest huddle for all the WTOxtr and WTWT lanes and evaluated the behaviors performed 15 mins prior to its initiation. We find that the duration of sniffs (anogenital investigation, see methods) by WT female choosers in WTWT lanes is greater than those by choosers in WTOxtr lanes (Fig. 4H). We did not find any difference in the duration of sniffs specifically with the winner or the loser when comparing WTWT and WTOxtr lanes, or in the total time spent sniffing over the course of the assay (Fig S4D, E). Thus, the interactions with both winner and loser contribute to the increase in duration spent sniffing 15 minutes prior to the longest huddle in WTWT lanes. These observations may suggest that a female chooser from the WTWT lane continues to seek chemosensory input before the longest huddle and these huddles are shorter due to ambivalence in a choice between two WT males. In contrast, a WT female chooser in a WTOxtr lane may encounter larger differences between a WT versus Oxtr^1-/-^ male, leading to a display of more prosocial behavior towards the chosen male independent of the winner’s Oxtr function.

We next examined antagonistic behaviors displayed during the naive choice assay as such behaviors may reduce prosocial behaviors displayed by chooser females. We found no difference in the overall counts of aggressive behaviors between WTWT and WTOxtr lanes (Fig. S4F). However, WT female choosers attack the loser male in the WTWT lanes more compared to WTOxtr lanes (Fig. 4I, Fig. S4G). Thus, in WTWT lanes, winner selection may be influenced by the level of aggression elicited by naïve WT males, a factor that is lower in WTOxtr lanes. We find a correlation between the amount of time spent socializing (with both winner and loser) and the count of all the aggression during the assay (Fig. 4J). Female choosers from WTOxtr lanes tend to spend more time engaged in prosocial behaviors with the winner (Fig. 4E-F, S4B) and display inceased prosocial and reduced aggressive behaviors. Female choosers choosing between two WT males, however, display either high prosocial and low aggressive behaviors or low prosocial behavior and high aggression. These observations suggest that the presence of a second male may influence the behavioral state of a WT female. The presence of only WT males may induce aggression by females against eventual losers, while such displays are absent in the presence of a male lacking Oxtr regardless of a female’s choice.

Given the differences in the influence of the presence Oxtr^1-/-^ males on displays of aggression by WT females when choosing between males, we examined antagonistic behaviors exhibited during interactions between naive (Introduction) or bonded (Stranger Rejection) females with WT or Oxtr^1-/-^ males in the absence of a third animal. Strikingly, we find that counts of aggressive behaviors were elevated only when naive WT females encountered WT males compared to Oxtr^1-/-^ males, while this difference is lost following pair bonding (Fig. 4K). In contrast, we find no differences in aggressive behaviors when comparing naïve or bonded WT and Oxtr^1-/-^ females (Fig. S4H). Taken together, these data suggest sex- and state-specific roles for Oxtr function. Specifically, in naive males, Oxtr function induces displays of aggression by naive females, and the presence of a male lacking Oxtr diminishes a naïve WT female’s propensity to display agonistic behaviors even in the presence of a WT male. Following bonding, however, the absence of Oxtr function in males does not appear to influence the behavior of WT females despite the promiscuous prosocial displays such mutant males.

Finally, to determine if naive prairie voles lacking Oxtr display generalized changes to their behaviors independent of attachment-associated social behavior, we examined general pro-social, anxiety-related, and locomotor behaviors. With the exception of an increase in locomotion displayed by Oxtr^1-/-^ females when compared to WT females in an open field paradigm, we found no differences between Oxtr^1-/-^ and WT voles of either sex (Fig. S4I).

We have examined social interactions during the initial contact between mates, the early stages of pair bonding, and during social attachment behaviors following partner preference formation. Consistent with our findings and previous work, Oxtr function is not required for the eventual display of partner preference. Here, we demonstrate that Oxtr function sex-specifically facilitates early social interactions that contribute to the formation of partner preference, and controls promiscuity.

### Oxtr controls the molecular signature of pair bonding in the nucleus accumbens

While partner preference formation can occur in the absence of Oxtr function, we find that it suppresses prosocial behaviors towards novel strangers following pair bonding. These results suggest that distinct components of pair bonding are differentially influenced by Oxtr function or compensated upon its loss. We next characterized the molecular signature associated with pair bonding to determine whether changes in gene expression following pair bond formation are controlled by Oxtr. Comparative studies demonstrate that Oxt binding in multiple brain regions differs significantly between promiscuous and monogamous rodent species. Oxt binding is particularly enriched in the NAc in prairie voles, while vasopressin binding is largely absent in this area^45,47^. Thus, analyzing gene expression in the NAc allows us to identify bonding-related expression that is primarily due to Oxtr function and largely independent of the direct consequences of AVP signaling. Additionally, given the well-established role of the NAc in the association of reward or aversion with specific sensory and environmental cues^69–71^ this region is well poised to play a role in the development of partner preference, even without Oxtr signaling, as well as stranger rejection, which appears to be dependent upon Oxtr function.

We performed bulk RNA-sequencing on NAc tissue from WT and Oxtr^1-/-^ animals of both sexes that were either: group housed with same-sex peers and sexually naïve (pre-pairing); or paired with WT opposite-sex partners for six days, using our timed mating paradigm (post-pairing) (Fig. 5A). PCA showed robust separation of WT and Oxtr^1-/-^samples (Fig. S5A). Differential expression (DE) analysis of pairing condition, regardless of genotype or sex, identified 13 genes that show significant changes in expression post-pairing (Fig. S5B), all of which showed decreased expression (Fig. 5B). These genes include *Agt, Sgk1, Ptgds, Fosb, Pdk4*, and *Acsm5* (Fig. 5C) and, while these genes have not previously been linked to shared gene regulatory or functional pathways, the predicted protein-protein interaction network for the 13 DE genes shows more connections than would be expected by chance (p=2.47 e^−05^) (Fig. S5C). Intriguingly, even in the limited set of genes with altered expression in the context of pair bonding, changes are driven primarily by expression in WT females (Fig. 5C, S5D). Subgroup analyses by genetic background and sex identified additional genes including *Adora2a, Fkbp5*, and *Spred3* in WT females and collagen-related genes, including *Col4a1* and *Col4a2*, in Oxtr^1-/-^ females (Fig. 5D, S5D). Only one gene, *Sema3b,* is significantly different in Oxtr^1-/-^ males following pairing (Fig. 5D).

**Figure 5:**
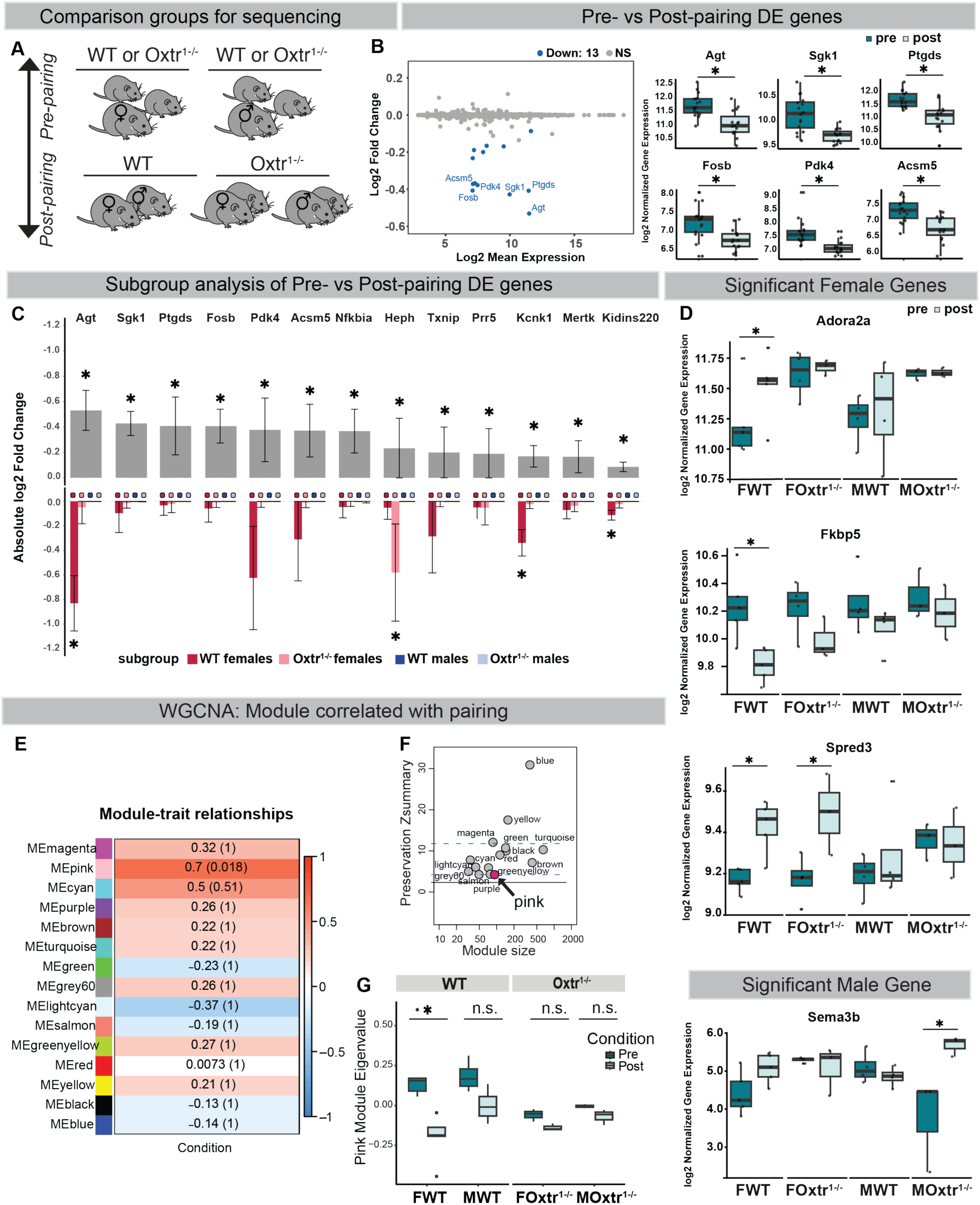
Post pair bonding gene expression is driven by female specific changes dependent on Oxtr signaling. A. Schematic of experimental groups used for sequencing highlighting pre- vs post-pairing comparisons between groups. Animals with symbols represent those used for tissue collection. B. Scatterplot illustrating the relationship between fold change and mean expression, emphasizing the 13 differentially expressed (DE) genes in post-pairing individuals compared to pre-pairing. Gray dots indicate non-significant genes (p_adj_ > 0.05); Blue dots indicate downregulated genes (p_adj_ < 0.05, log_2_FC <0) and upregulated genes in red (p_adj_ < 0.05, log_2_FC>0). The right panel shows box plots of normalized gene expression for the top six DE genes by fold-change between pre- and post-pairing conditions. C. Bar graph emphasizing how each combination of genotype and sex (denoted at bottom as ‘subgroup’) influence the 13 overall pairing-DE genes identified. The top graph depicts the absolute fold change and standard error of the differential expression analysis including all pre- and post-bonding individuals in which the 13 genes of the x axis were identified. The bottom graph depicts the absolute fold change and standard error in analyses when only individuals of that subgroup were considered. (* denotes p_adj_ < 0.05 for the pre- vs post-pairing analysis). D. Box plots of select genes that show sex- and genotype-specific changes in expression with pairing condition. *Adora2A* and *Fkbp5* are DE when considering WT females, *Spred3* is DE when considering WT and Oxtr^1-/-^ females, and *Sema3b* is DE when considering Oxtr^1-/-^ males. Note: Sema3b is the mouse ortholog gene symbol for vole gene “ENSMOCG00000019704” which does not have an assigned gene symbol. E. Heatmap showing correlations between module eigengenes (MEs) for modules^WT^ and pairing condition (Pre=1, Post=0). Heatmap color and value reflect Pearson correlation coefficient; adjusted p values shown in parentheses (p_adj_ = p * number of modules). F. Preservation of modules^WT^ in OXTR-mutant samples. The Z_summary_ measure combines module density and intramodular connectivity metrics to a composite statistic where Z>2 suggests moderate preservation and Z>10 suggests high preservation^134^. Of all modules^WT^(n=15), Mpink^WT^ had the lowest Z_summary_ (Z_summary_=2) in mutant samples. G. Box plots showing comparison of MEpink^WT^ for pre- vs post-pairing condition separated by sex and genotype. (Wilcoxon rank sum test, Bonferroni adjusted for 4 comparisons). For all box plots center = median, box boundaries = 1^st^ and 3^rd^ quartile, whiskers = 1.5*IQR from boundaries. (*n* = 16 pre-pairing and 15 post-pairing) See Supplementary Figure 5.

To further examine changes in patterns of gene expression in the NAc associated with pair bonding, we performed weighted gene coexpression network analysis (WGCNA)^72^ and calculated three sets of modules (M) from different sample subsets (modules^WT^, modules^Mut^, and modules^All^) (Fig. S5F, S6D). Using the modules^WT^ subset, we identified 15 coexpression modules (Fig. S5F, Table 2) representing genes that share highly similar expression patterns within the NAc. We identified one module (Mpink^WT^) that significantly and specifically correlated with pre-pairing status in WT animals (Pearson R correlation of 0.7, Bonferroni adjusted p-value (p_adj_)=0.018) (Fig. 5E, S5E-F). Genes with high module membership for the Mpink^WT^ are also those with high gene significance for pairing condition supporting the module’s association with behavior condition (Fig. 5SG). Further, genes significantly differentially expressed with bonding are significantly enriched in Mpink^WT^ (Bonferroni p_adj_=7.44 e^−08^) and serve as hub genes within this module (Fig. S5E, G). These results suggest that the Mpink^WT^ module might be driving, or under the influence of pathways driving, the pre-pairing to post-pairing state change.

We asked whether any of the modules^WT^ identified were regulated by Oxtr and thus lost in Oxtr null animals. To address this, we assessed how well modules^WT^ are preserved in mutant samples. We calculated a Z_summary_ measure that combines module density and intramodular connectivity metrics into a composite statistic. We find that of the modules^WT^, Mpink^WT^ had the lowest Z_summary_ (Z_summary_ = 2) in Oxtr^1-/-^ samples (Fig. 5F), indicating that dynamics of gene expression that correlate with bonding states are lost in the absence of Oxtr. Finally, the module eigengene (ME) for Mpink^WT^ when calculated for all 30 samples and compared between pre- and post-pairing, was significant only for WT females (Wilcoxon rank sum, Benjamini-Hochberg p_adj_=0.03) and not for WT males or Oxtr^1-/-^animals of either sex (Fig. 5G). These observations suggest there is sex-specific, coordinated regulation of gene expression in the NAc following pairing of mates that may be associated with pair bond associated behaviors.

### Loss of Oxtr sex-specifically disrupts patterns of gene expression associated with neurodevelopmental disorders

We next wished to determine how Oxtr signaling contributes to changes in gene expression in the NAc associated with the naive state or the process of pair bonding (Fig. 6A) by comparing the transcriptional profile of WT and Oxtr^1-/-^ animals of each sex. Given that our lines are maintained on an outbred background, in order to minimize sample variability, tissue from three animals was pooled for each sample. We observed significant genotype specific differences in gene expression in animals from both sexes, with 466 genes showing differences between WT and Oxtr^1-/-^ females, 270 between WT and Oxtr^1-/-^ males, and 1,014 genes when samples from both sexes were combined (using a p_adj_<0.05 and absolute (log_2_(FC)) > 0.25 as criteria). Of the combined DE gene set, 370 genes are upregulated and 644 are downregulated (Fig. 6B, S6A-B). Genes with the greatest fold change across both sexes include *Slc2a3, Kcna1, Rgs8, Cnksr2, Acvr*, and *Cnot6* (Fig. 6B). Genes specifically downregulated in Oxtr^1-/-^ females with log_2_(FC) less than 1 include *Abcb11* (a gene recently associated with treatment resistant schizophrenia^73^), *Ngfr, Pon3,* and *Shh*. While there was no significant difference in Oxtr transcript levels by genotype (p_adj_ = 0.753)—consistent with minimal nonsense-mediated decay despite a premature stop codon—we have previously shown that no functional protein is produced in Oxtr^1-/-^ animals via autoradiography^59^. V1aR showed a modest decrease in expression in Oxtr^1-/-^ animals (p_adj_=0.03) (Fig. S6C), though this is likely influenced by variability in WT samples. WGCNA of all samples identified two modules (Mbrown^All^ and Mturquoise^All^) that were significantly positively correlated with the WT genotype compared to mutant (Pearson R = 0.55, p_adj_ = 0.011 for brown and Pearson R = 0.8, p_adj_ = 6.6e-7 for turquoise) (Fig. S6D). None of the identified modules showed correlation with pairing condition (Fig. S6D). Our results thus reveal a dramatic change in patterns of gene expression in the NAc with global loss of Oxtr signaling, independent of bonding state.

**Figure 6:**
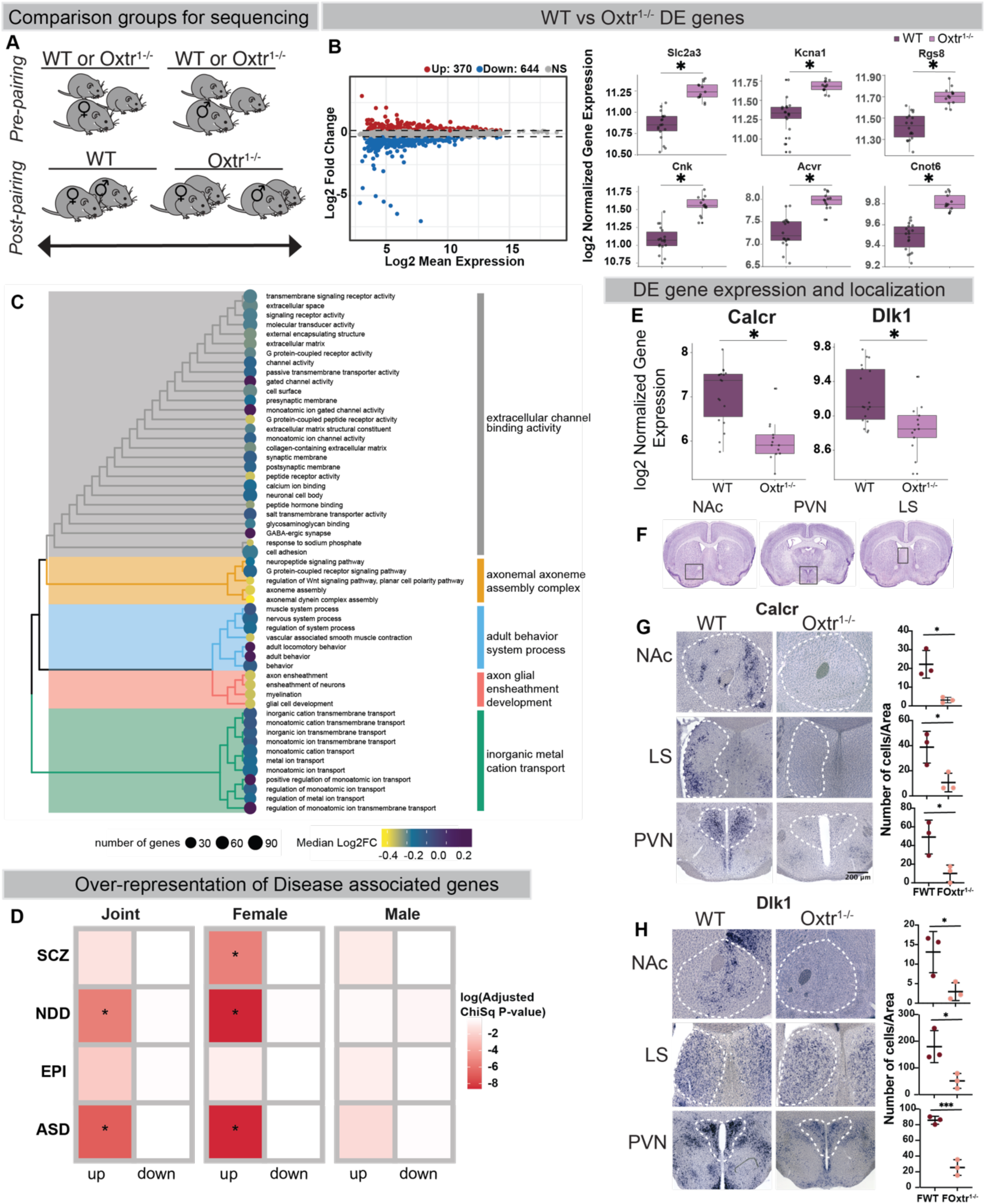
Oxtr regulated genes are common to social behavior relevant processes across species and are similarly regulated across brain regions. A. Schematic of experimental groups for sequencing, highlighting genotype comparisons between groups. Animals with symbols represent those used for tissue collection. B. Scatterplot illustrating the relationship between fold change and mean expression, emphasizing differentially expressed (DE) genes in Oxtr^1-/-^ individuals compared to WT. Significantly downregulated genes are shown in blue (log_2_FC<-0.25, p_adj_<0.05) and upregulated genes in red (log_2_FC>0.25, p_adj_ < 0.05). Box plots of normalized gene expression for the top six DE genes by fold change between genotypes. C. Gene Ontology enrichment analysis for all genotype-DE genes (n=1014) highlights processes related to extracellular channel binding activity, axonal processes, behavior related gene expression, and cation transport. The leaves represent significant GO categories (BH adjusted p value < 0.05) which have been hierarchically clustered by their semantic similarity. Branches are colored by cluster and high-frequency words are displayed to the right. Leaf size and color correspond to the number and median fold change of DE genes in each category. D. Enrichment analysis shows that genes which are DE in Oxtr^1-/-^ samples are over-represented in some neuropsychiatric disease-associated gene sets. The first DE gene set evaluated (‘Joint’) represents genes which are DE in Oxtr^1-/-^ samples when considering both sexes in the differential expression analysis (p_adj_ < 0.01). The second gene set are DE genes when only considering females, and the third when only considering males. Gene sets are further separated by direction of effect across the x-axis to highlight the enrichment in upregulated DE genes. Colors correspond to the Benjamini-Hochberg corrected Chi-square p values; asterisks denote p_adj_ < 0.05. Y-axis abbreviations are as follows: SCZ=schizophrenia, NDD=neurodevelopmental disorder, EPI=epilepsy, and ASD=autism spectrum disorder. E. Box plots of *Calcr* and *dlk1* normalized gene counts by genotype. Log_2_FC:-0.962 and p_adj_: 1.75e-06 for *Calcr* and log_2_FC: −0.276 and p_adj_: 0.00334 for *dlk1* from comparison considering all WT and Oxtr^1-/-^ samples. F. Brain regions selected for in situ analysis. (NAc=nucleus accumbens, PVN=paraventricular nucleus of the hypothalamus, LS=lateral septum) G. ISH images and quantification for *Calcr* expression in NAc, PVN, and LS from WT and Oxtr^1-/-^ animals. H. ISH images and quantification for *Dlk-1* expression in NAc, PVN, and LS from WT and Oxtr^1-/-^ animals. Box plots in G. and H. show comparison of the area-normalized cell number by genotype. (*n*=3 per condition, Wilcoxon sign rank, *=p<0.05, ***=p<0.001) See Supplementary Figure 6 and Supplementary Table 1.

Examination of the sets of genes whose expression is altered by the loss of Oxtr, regardless of bonding state or sex, revealed several distinct cellular processes regulated by Oxtr signaling. These processes comprise larger domains of cell function including extracellular channel binding activity, axoneme assembly complex, axon glial ensheathment development, and metal cation transport (Fig. 6C), as well as pathways for G protein signaling and dopaminergic neurogenesis (Fig. S6E). When analyzed by sex, more of the identified processes were enriched in female Oxtr gene sets than in male ones (female: 19/62, male: 6/62). To further link differentially expressed genes to behavioral traits, we asked whether genes altered by loss of Oxtr are selectively enriched for high-confidence associations with autism spectrum disorders (ASD) and other neurodevelopmental diseases (NDD) affecting social attachment in humans as compared to epilepsy, a disorder that does not disrupt social attachment^74–77^. Consistently, we find that the genes upregulated with loss of Oxtr (p_adj_<0.01, log_2_(FC)>0) are specifically enriched for ASD- and NDD-associations but not epilepsy associations (Fig. 6D “Joint”). This enrichment is again primarily driven by genes upregulated in females and not males (Fig. 6D). Thus, similar to the factors that influence gene expression changes associated with pair bonding, many of the changes we identify in the NAc resulting from loss of Oxtr are primarily driven by female-specific effects that may play a conserved role in mediating social attachment in humans.

### Oxtr regulates coordinated gene expression across multiple neural populations implicated in social attachment behaviors

Given the expression of Oxtr in regions of the brain known to mediate social behavior in voles and other species^78,79^, we wished to determine if genes whose expression is altered in the NAc by the loss of Oxtr were expressed not only in specific populations of cells within this region, but in other regions of the prairie vole brain implicated in social and attachment behaviors. We therefore examined the expression of 15 genes that showed significant changes in expression in WT females when compared to those lacking Oxtr using *in situ* hybridization (ISH) to both validate our RNA-seq findings as well as examine the expression patterns of these genes. Several of these genes, such as phosphodiesterase 10A (*Pde10A)* and diacylglycerol kinase beta (*Dgkb)*, show robust expression throughout the striatum (data not shown), while genes such as the calcitonin receptor (*Calcr*) and delta like non-canonical notch ligand 1 (*Dlk-1*) are expressed in restricted populations of cells in the NAc (Fig. 6F-H, top panels). The expression of these genes is significantly reduced in Oxtr^1-/-^ females (Fig. 6E-H), supporting the specificity and resolution of our RNA-seq findings.

In parallel to our examination of gene expression in the NAc, our ISH revealed that loss of Oxtr in females altered the expression of the genes we examined in other regions of the prairie vole brain. We examined the expression of these genes in the lateral septum (LS), a region that has been implicated in agonistic and attachment behaviors in prairie voles and other rodents and shows species dimorphic patterns of Oxt and Avp binding. We observe that both *Calcr* and *Dlk-1* are decreased in the absence of Oxtr (Fig. 6G, H middle panel) in the LS. We also find that expression of *Calcr* and *Dlk-1* is decreased in the paraventricular nucleus of the hypothalamus (PVN), a principal source of Oxt and Avp in the rodent brain which sends projections to the NAc (Fig. 6G-H bottom panels)^80–82^. These findings suggest that Oxtr function regulates patterns of gene expression in multiple regions of the brain implicated in social and attachment behaviors in a coordinated and region-specific manner.

### Loss of Oxtr results in the decrease in density of Oxt and Avp expressing neurons in the PVN

We find here that loss of Oxtr results in coordinated changes in gene expression in multiple neural populations, including the PVN. We therefore wished to determine if the signatures of gene expression regulated by <u>O</u>xtr in the <u>N</u>Ac in <u>p</u>rairie voles (ONP set), regardless of sex and bonding state, were enriched for patterns of gene expression associated with distinct neural populations in other regions of the brain, particularly those implicated in social and attachment behaviors. We compared the ONP gene set to the molecular signatures of 33 hypothalamic subtypes previously identified by single-cell RNA sequencing in mice^83^. Of the 33 clusters examined, the four with highest overlap with the ONP gene set represent subsets of PVN neurons, in particular those that express Oxt and AVP (Fig. 7A, clusters 43 and 26), as well as PVN neurons that express *Dlk1* and *Calcr* (cluster 29), both of which show reduced expression in the NAc and PVN in females lacking Oxtr (Fig. 6G-H bottom panel).

**Figure 7:**
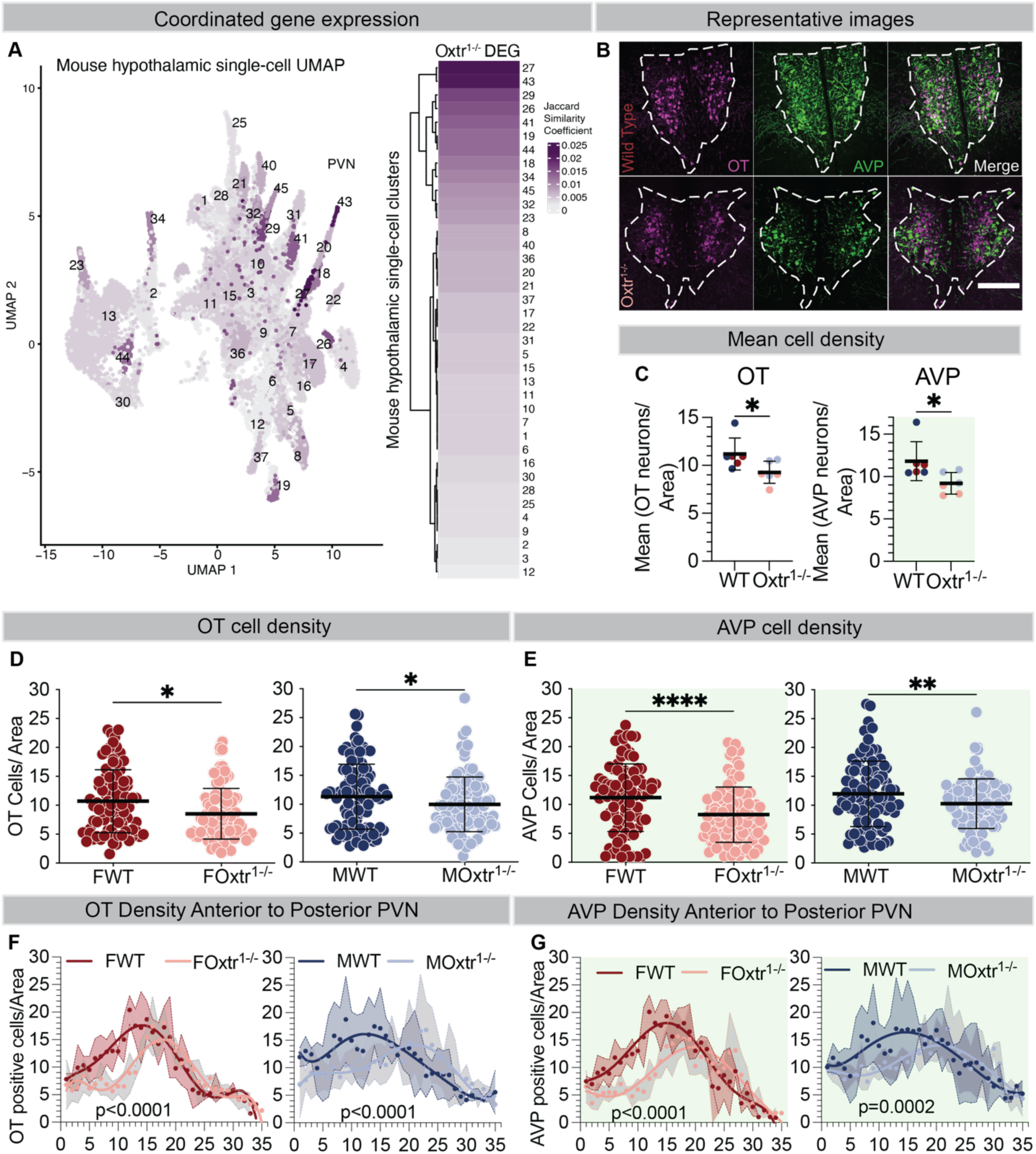
Loss of Oxtr results in the decrease in density of Oxt and Avp expressing neurons in the PVN. A. UMAP visualization of mouse hypothalamus single-cell RNAseq dataset^83^ with proposed PVN clusters labeled. The authors’ original numeric cluster labels and embedding are shown. Each cluster is colored by the Jaccard similarity coefficient between the mouse marker gene sets and the vole Oxtr^1-/-^ DE genes (ONP set). This is also represented in a heat plot to the right. Some of the highest similarity is seen in Cluster 43, which represents Oxt+ cells, and Cluster 26, which represents Avp+ cells in the PVN. B. Representative images of IHC staining against OT and AVP neuropeptides from FWT and FOxtr^1-/-^ PVN showing the loss of structural integrity and OT and AVP positive cells from the knock-out. (White dotted line denotes the PVN area, scale bar represents 100µm) C. Mean of positive cells per unit area across all sections is lower for Oxtr^1-/-^s vs WT for both OT (left) and AVP (right). Each dot represents 1 animal. D. The number of sections with a high density of OT positive cells is greater in both female (left) and male (right) WT animals. E. Similarly, WT animals of both sexes show an increase in the number of sections containing AVP positive neurons in high densities. F. The average OT cell density from matched PVN sections arranged from anterior to posterior shows that the decrease in cell density in Oxtr^1-/-^ animals is biased towards the anterior. Each dot represents the average cell density, the lines represent the sixth order non-linear regression model for cell density by sex and by genotype (n=3) and the ribbons represent the standard deviation. G. Oxtr^1-/-^ animals show a significant decrease in AVP cell density in the anterior PVN in both sexes. See Supplementary Figure 7 and Supplementary Table 1. Mean +/− SD, n = 3, *p =< 0.05, FWT = female WT, FOxtr^1-/-^ = female Oxtr^1-/-^, MWT = male WT, MOxtr^1-/-^ = male Oxtr^1-/-^.

We next sought to determine if the overlap between Oxtr regulated genes and gene expression patterns of neural populations in the PVN correlated with changes in the expression of neuropeptides within PVN itself. We examined patterns of Oxt and Avp expression using immunohistochemistry in naive adults and compared patterns of expression between WT animals of both sexes and those lacking Oxtr (Fig. 7B, Fig. S7A-B). We find that the density of both Oxt and Avp producing neurons is lower in Oxtr^1-/-^ animals (Fig. 7C-E). To determine if the loss of Oxt or Avp expression was restricted to specific populations of PVN neurons, we examined the expression of these peptides across the anterior-posterior axis of the PVN in both WT and Oxtr^1-/-^ males and females. We find that animals of both sexes show a preferential loss of Oxt and Avp expressing neurons in the anterior PVN (Fig. 7F-G). Finally, we examined patterns of Oxt and Avp over the course of development to determine when loss of Oxtr function influences the development of their expression. We find that the density of Oxt neurons at P0 (Fig. S7C-F) and both Oxt and Avp neurons at P21 is lower in Oxtr^1-/-^ females compared to age matched WT females (Fig. S7G-J). In contrast, we find that Oxt and Avp neuron density loss occurs only in adult Oxtr^1-/-^ males (Fig. 7D-G, Fig. S7B). These observations suggest the coordinated, region-, and sex-specific influence of Oxtr function on the development of neurons expressing Oxt (and Avp) within the PVN, consistent with a trophic role for this pathway identified in mice^63,65,84,85^.

## Discussion

Robust reproductive behaviors are essential for survival. Accordingly, species have evolved diverse strategies, including social monogamy, which mediate locating, identifying, and attracting mates, facilitate reproduction, and promote the survival of progeny^16,86^. Signaling via Oxtr influences a wide range of physiology and behavior associated with social interactions and attachment between conspecifics, including pair bonding between prairie vole mates^41,87^. Acute blockade of Oxtr signaling via central or nucleus accumbens infusions of Oxtr antagonists prevents the formation of partner preference^28,50,51,60,88^. In contrast, constitutive loss of Oxtr in prairie voles disrupts specific components of pair bonding, while the capacity for partner preference ultimately remains intact^59^. Similarly, Oxtr null mice show select behavioral deficits, including impaired sociability, reduced preference for social novelty, and increased aggression towards a novel intruder^89,90^.Thus, like other circuit and neuromodulatory pathways across diverse species, Oxtr likely functions in parallel to multiple genetically specified pathways to influence specific behavioral modules that comprise complex behavioral states, including social attachment.^33,35,91–97^. These actions are likely dependent on the developmental context and engagement of potentially compensatory parallel pathways, leading to divergent phenotypes depending on the timing and nature of Oxtr disruption.

Here we demonstrate that loss of Oxtr in prairie voles influences the dynamics of social interactions between sexually naïve animals and WT potential mates in distinct social contexts in sex-specific ways. The presence of a naive Oxtr^1-/-^ male changes levels and patterns of both early prosocial and antagonistic behaviors displayed by naive WT females, supporting the role of Oxtr in controlling individuals’ and reciprocal behavior ^30^. Thus, Oxtr function orchestrates male social behaviors operating early in the sensitive period of pair-bond formation and, through these behaviors or through additional sensory cues yet to be identified (pheromone signatures or vocal communication), shapes bonding.

Animals lacking Oxtr take longer to display partner preference, and despite strong partner preference following longer cohabitation, demonstrate promiscuous prosocial behaviors towards novel potential mates, including huddling with strangers. These observations are consistent with a state-dependent role of Oxtr function, similar to findings in multiple species, including humans^98–100^. Oxtr function may thus promote prosocial behavior in the naïve state to facilitate attachment with an unfamiliar potential mate, but following pair bond formation, suppresses further prosocial interactions to new individuals, or even promotes further separation from and rejection of unfamiliar individuals or groups. Across species and outside of the context of pair bonding, oxytocin signaling may play a conserved role in mediating other state-specific aversive behavior, for example, defeat-induced social avoidance learning^101^.

Our findings reveal that patterns of social behaviors contributing to 1) pair bond formation, 2) partner preference, and 3) rejection of novel mates are genetically separable components of pair bonding representing distinct behavioral modules regulated by mechanisms that require Oxtr function, i.e. reciprocal behaviors and stranger rejection, or those that can be displayed in its absence, i.e. partner preference. Further work will allow us to determine the Oxtr-independent mechanisms that mediate the closure of an early sensitive period and facilitate the display of partner preference^59^. Where Oxtr functions to control the switch from prosocial behavior towards a potential mate in naïve animals to the suppression of such behaviors towards, and active rejection of, opposite-sex conspecific strangers following bonding remains to be determined.

Examination of the molecular signatures of pair bonding and the role of Oxtr in regulating these changes in gene expression in the NAc reveals modest, Oxtr-dependent changes in the expression of a specific network of genes in WT animals following pair bonding^45,102^. Intriguingly, loss of Oxtr preferentially impacts patterns of gene expression in the NAc in females. The NAc demonstrates little endogenous sex steroid hormone receptor expression difference in gene expression in prairie voles and other species^78,103–105^. Sex differences in the effects of Oxtr on gene expression this region may therefore suggest that there are significant sex-differences in the inputs into this region that act through Oxtr during development^106–111^. The larger impact of loss of Oxtr in females is consistent with its more general conservation in mammals to regulate aspects of female physiology such as lactation and prosocial behaviors or sensory cues that promote interactions with males^112^.

Oxt and its receptor show species-specific changes in expression through embryonic development and early postnatal life, impacting the development of cortical and subcortical circuits^62,64,113,114^. Consistent with this, genes with altered expression in the absence of Oxtr are enriched for genes associated with neuropsychiatric disorders, in particular neurodevelopmental disorders and ASD. Given the sex-biased enrichment of these genes, such developmental differences in the impacts of Oxtr function may contribute to sex differences in the pathophysiology of these disorders^115–117^. Accordingly, the molecular signatures of attachment in the NAc, both dependent upon and independent of Oxtr function, are subtle relative to these sex-biased differences in the role of Oxtr and possibly reside in other neural populations that function in attachment.

Validation of changes in gene expression in the NAc resulting from loss of Oxtr suggests coordinated regulation of these genes by Oxtr across multiple brain regions implicated in social and attachment behaviors. Oxtr may be one of multiple factors that organizes specific components of the circuits underlying these displays to control discreet modules of behavior. Consistent with a trophic role for Oxtr function during development, we find that the genes regulated by Oxtr within the NAc are enriched for those that identify cell types in the PVN, a principal source of neuropeptides implicated in pair bonding, which sends oxytocinergic projections to the NAc^79,81,118–120^. Neonatal OT manipulations in mice sex-specifically modulate levels of neuropeptides in PVN neurons, increasing Oxt expression in females and reducing Avp expression in males^64^. We demonstrate that, in prairie voles, loss of Oxtr preferentially impacts Oxt and Avp expression in the anterior PVN, suggesting that specific populations of these cells may be more dependent upon such trophic function for their development, survival, or expression of these hormones. Whole brain analysis will likely reveal if such neurons have distinct projections when compared to those that remain in the absence of Oxtr, contributing to our understanding of how these circuits and pathways regulate distinct aspects of social and attachment behaviors and physiology^4,87,121^.

Taken together, our behavioral and molecular studies reveal previously unappreciated nuances in the control of social behavior and pair bonding by Oxtr function. They uncover intriguing sex differences in the patterns of behavior influenced by loss of Oxtr function. These observations suggest that circuits with sex-specific patterns of development, or even function in ancestral species, may mechanistically converge in the context of Oxtr function to generate displays of reciprocal interactions, pair bonding and attachment behaviors in WT prairie voles that are similar between males and females^122,123^. Manipulation of Oxtr expression at specific developmental timepoints and in specific neural populations will help to elucidate how this pathway influences social and attachment behaviors in the context of normal development to regulate distinct components of pair bonding. Understanding the specific neural populations that express Oxtr and other molecular pathways that contribute to pair bonding will allow us to understand the mechanisms controlling the early experience-dependent, sensitive period of bond formation. In parallel, examination of the activity in these populations following manipulations of Oxtr function will allow us to determine the circuit mechanisms by which similar but distinct sensory cues contribute to the identification of partners or novel conspecifics and consequently, the display of dramatically different patterns of social behaviors. The sum of these mechanisms represents the preference for partners over strangers and reflects the formation and maintenance of pair bonds between mates, affiliation between individuals, and the behavioral consequences of identifying others as unfamiliar or outside of these bonds.

## Acknowledgments

The authors would like to thank members of the Manoli lab for assistance, advice, helpful discussion and comments on the manuscript. J. Tollkuhn and M. Brainard provided valuable insights to the structure, content and design of the manuscript. Additionally, we would like to thank S. Garg for his assistance in data analysis and help designing the analytic pipeline. The authors received funding from National Institutes of Health grant R01MH123513 (D.S.M.), National Science Foundation grant 1556974 (D.S.M.), Burroughs Wellcome Fund 1015667 (D.S.M.), Whitehall Foundation grant 2018-08-83 (D.S.M.), One Mind Foundation A137726 (D.S.M.), National Institutes of Health grant R25MH060482 (K.M.B.), AP Giannini Foundation Fellowship P0534952 (K.M.B.), Larry L. Hillblom Foundation Fellowship 2020-A-023-FEL (K.M.B.), and National Institute of Mental Health grant R01MH123178 (K.S.P.)

## Author contributions

Conceptualization, R.S., K.M.B., K.L., and D.S.M.; methodology, A.B., A.J.W., D.S.M., K.L.B., K.M.B., K.S.P., and R.S.; software, K.Q., and S.W.; formal analysis, A.E., R.S., and K.M.B.; investigation, R.S., K.M.B., A.E., B.W., G.W., S.W., K.Q., R.D.L., K.L., N.H., B.A.S., M.C.H., M.S., R.K., A.C., D.G., L.C.C., N.L.G.; visualization, R.S., K.M.B., B.W., and A.E.; funding acquisition, D.S.M., K.S.P., and A.J.W; writing– original draft, R.S., and K.M.B.; writing-review & editing, R.S., K.M.B., D.S.M., N.H., K.S.P., A.E., and A.B.

## Star Methods

### Key resources table

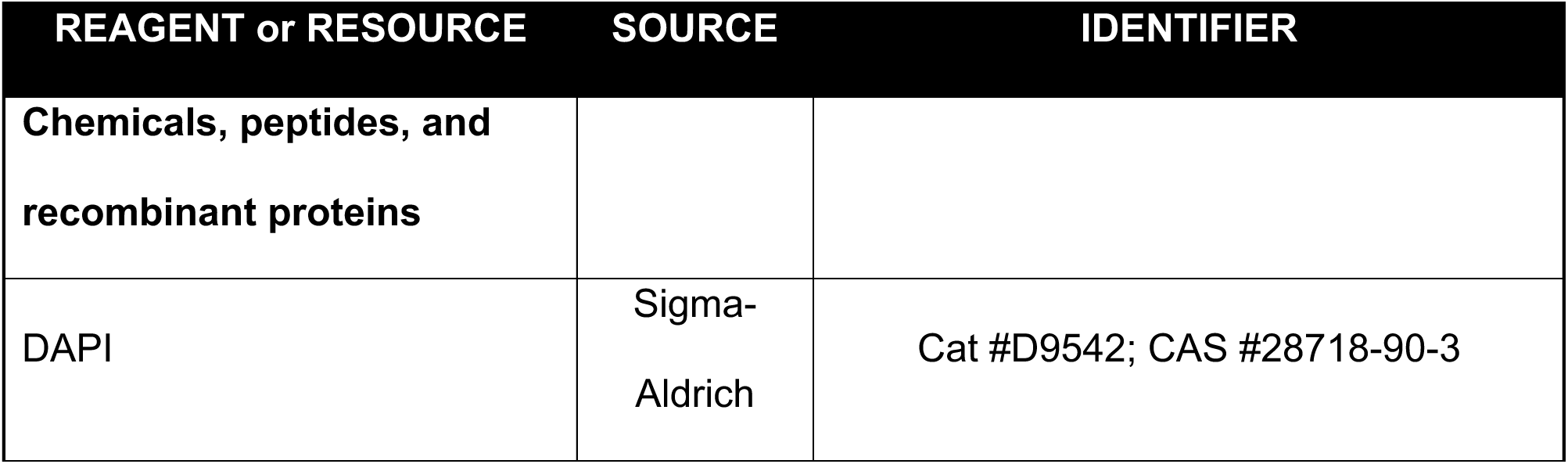
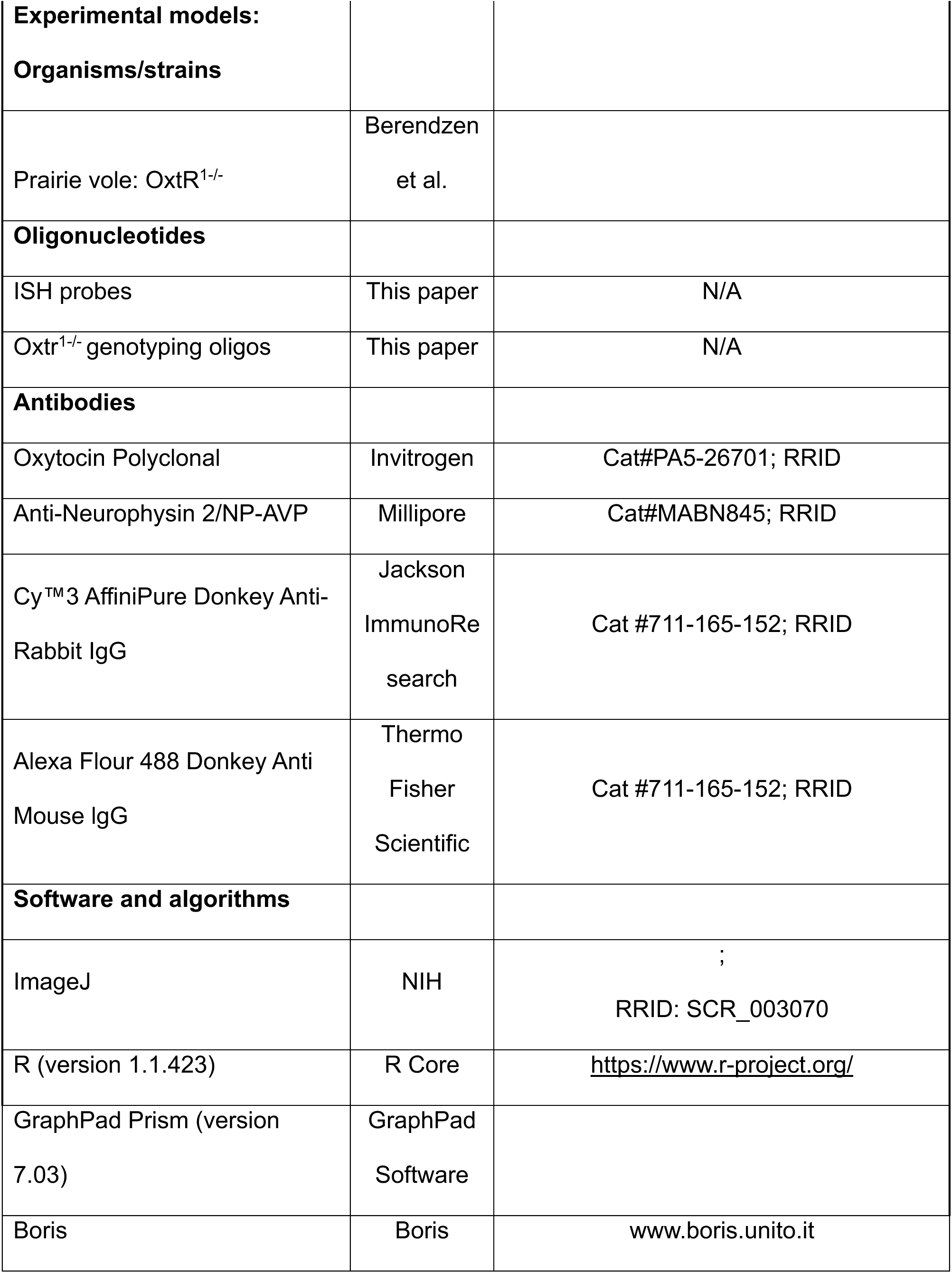
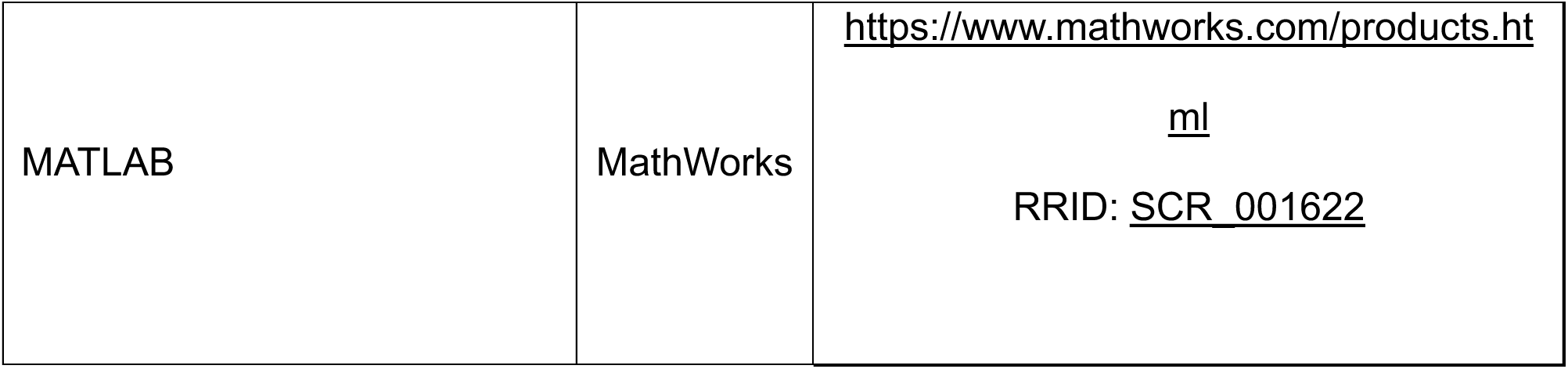

### Resource availability

#### Lead contact

Further information and requests for resources and reagents should be directed to and will be fulfilled by the lead contact, Devanand Manoli.

#### Materials availability

All unique reagents generated in this study are available from the lead contact upon request.

#### Data

Transcriptomics data discussed in this publication are accessible through GEO Series accession number GSE279248.

#### Code

All code used for RNA-seq data processing and differential expression analyses, as well as post-processed expression data, can be found at https://github.com/aseveritt/OXTR_prairie_voles

### EXPERIMENTAL MODEL AND SUBJECT DETAILS

#### Animals

Subjects were laboratory-bred prairie voles (*Microtus ochrogaster*) which originated through systematic outbreeding of a wild stock captured near Champaign, Illinois. CRISPR/Cas9 editing was used to generate 2 independent null alleles, Oxtr^1-/-^ and Oxtr^2-/-^, both of which were validated using autoradiography to show loss of receptor function^59^ (data not shown). Oxtr^1-/-^ was backcrossed for >7 generations to mitigate any possible off-target effects and this line was used for all subsequent experiments. Sexually naïve male and female animals were group weaned at 21 ± 1 days and separated to group housing with same-sex siblings and age-matched same-sex non-siblings. Voles were maintained under a 14:10 h light-dark cycle in clear plastic cages (45 × 25 × 15 cm) with bedding, nesting material (nestlet), and a PVC hiding tube. Rooms were maintained at approximately 20°C, and food and water were available *ad libitum*.

Breeding pairs were established between two heterozygotes or a homozygous Oxtr^1-/-^mutant male and heterozygous female partner from a breeding line maintained in the lab. Voles were randomly assigned into experimental groups when they reached 7-9 weeks of age at the start of testing. This study was carried out in accordance with the recommendations in the Guide for the Care and Use of Laboratory Animals published by the National Research Council. The protocol was approved by the Institutional Animal Care and Use Committee at each respective institution.

#### Genotyping

The following primers were used to amplify a region where XcmI restriction for Oxtr^1^ or Bpu10I restriction for Oxtr^2^ determined the genotype of all animals studied. XcmI cuts the mutant allele in the case of Oxtr^1^ and Bpu10I restricts the WT sequence for Oxtr^2^. (Forward: ACTGGAGCTTCGAGTTGGAC, Reverse: ATGCCCACCACTTGCAAGTA)

### METHOD DETAILS

#### Behavior

##### Triadic naive choice assay

The linear three chamber partner preference apparatus with open top and 10 x 32 inch walls^59^ was modified to capture the interactions of the naive, freely moving WT subject and the naive, tethered, opposite sex potential partner (either 2 WTs or a WT and an Oxtr^1-/-^) at either end of the chamber. Behaviors were recorded for 6 hours using a top view camera and the videos generated were analyzed using a contour detection algorithm. Specifically, contours were detected on individual video frames using a threshold-based strategy that took advantage of the contrast between dark-colored vole fur coats and light-colored bedding and chamber apparatus. A standalone vole was detected as a single contour, and two voles in physical contact were likewise detected as a single contour. Consistent vole identities across video frames were determined using the Hungarian algorithm. Since the assay began with one vole in each of the three chambers, and since the subject vole remained freely mobile throughout, it was possible to unambiguously assign each contour to a single vole identity (e.g., subject, right chamber, or left chamber) or to that of a pair of voles (e.g., subject and right chamber voles or subject and left chamber vole), resulting in continuous trajectories for each of the three animals. Time in chamber was determined by the chamber location of the subject vole in each frame, since the chamber location of the right chamber and left chamber animals was fixed throughout the assay by physical tethering and any time the contour of the subject animal merged with that of the tethered animal (right or left), the subject was termed in contact with it. The algorithm also determined the boundaries of the chambers and calculated chamber times. We used the chamber time data from when both ends of the chamber contained tethered WTs (WT-WT condition) to ensure that there was no bias in which side the choosers preferred. This footage was then scored manually to produce greater resolution of behaviors.

##### Bond formation assays

Initial introduction of Oxtr^1-/-^ and WT animals of either sex with an opposite sex WT animal was filmed for 30 minutes. Observed behaviors included anogenital sniffing, investigation (subject animal’s nose contacts any non-genital region), duration of stationary huddling or >50% side-by-side contact between animals, rearing, frequency of aggressive behavior including lunges, receipt of aggression, and tussles, as well as mating behavior including mounting, intromission, and ejaculation. These animals remained in their home cage for 18 hours following pairing, after which, they were separated by a clear, plastic barrier. The barrier was left in place for 24 hours and then removed allowing the subject free access to the partner animal thereafter. A 30-minute period directly following barrier removal was also filmed to assess mating behavior, termed the timed mating assay.

Subjects remained cohoused with the opposite-sex, wildtype partner for durations as indicated in the text. The partner preference test (PPT), as previously published^59^, was used to assess pair bond formation. Subject, partner, and an opposite sex stranger are placed in a three-chamber apparatus with open top and 10 x 32-inch walls. The partner and stranger animals are tethered on either end of the three-chamber arena and the subject allowed free access over a period of 3 hours. Behaviors are recorded from a top view camera capturing the entire apparatus. Videos were scored using BORIS^124^ post-test by validated scorers blind to condition. Observed behaviors included location (i.e., duration of time in partner, stranger, and neutral chambers), duration of stationary huddling or >50% side-by-side contact between the partner and stranger animals, and frequency of aggressive behavior (i.e., lunges). Of note, analysis of partner preference between wild type siblings from the Oxtr^1-/-^ background and wild type animals from the breeding colony showed no difference in time spent huddling with partner vs stranger animals.

##### Bond maintenance assays

Four days after the timed mating assay, female animals underwent a separation-reunification assay. Females remained in their home cage while their male partners were placed in a separate cage in another room for one hour. The males were then reintroduced into the home cage and behavior was video recorded for 30 minutes. Male subject animals underwent the same procedure five days following timed mating. Two days following the separation reunification test, animals underwent a selective aggression assay. Partners were again removed from the home cage and separated to a separate room for an hour. After one hour, a novel sexually naïve animal of the opposite sex was introduced into the home cage with the subject animal and behavior observed for 20 minutes. If significant aggression occurred, defined as >3 tussles during the assay, the animals were separated and the assay terminated. Overall, two WT female assays and one Oxtr^1-/-^ were terminated for aggression. The same behaviors listed above for the introduction were observed and recorded in the reunification and aggression assays.

#### RNA Sequencing

##### Sample collection

Adult 7-9 week old WT and Oxtr^1-/-^ voles were either cohoused with same sex-siblings or paired with an animal of the opposite sex for four days following the timed mating protocol described above. These animals were euthanized by CO_2_ inhalation and decapitated. The brain was quickly dissected out and sectioned into 500μm coronal slices using a brain matrix mold (BrainTree Scientific) chilled on ice. Slices were floated in ice-cold phosphate buffered saline (PBS), the NAc was identified using anatomical landmarks, and dissected using a Zeiss microscope. A total of 45 WT females, 39 WT males, 21 Oxtr^1-/-^ females, and 18 Oxtr^1-/-^ males were split between paired and unpaired conditions and tissue from three animals per condition was pooled prior to RNA extraction. Tissue was flash-frozen in liquid nitrogen and stored at −80°C until further processing. Total RNA was extracted using TRIzol according to manufacturer’s instructions and quantified using NanoDrop (Thermo Scientific). Library preparation for next generation sequencing, comprising 35 samples and 4 technical replicates, was done using TruSeq Stranded mRNA Kit according to manufacturer recommendations and sequenced on a NovaSeq 6000 to an average depth of 2.9e07 reads per sample. Sequencing was performed across three runs, but all data was processed together.

##### Read processing

Adapter and low-quality sequence trimming was performed with Trimmomatic v0.39 using parameters: ‘TruSeq3-PE.fa:2:30:10:2:keepBothReads LEADING:3 TRAILING:3 SLIDINGWINDOW:4:20 MINLEN:25’ (Bolger et al., 2014). Trimmed RNA-Seq reads were aligned to a custom MicOch1.0 prairie vole genome, which incorporated a known missing gene V1A, using STAR v2.7.3a in gene annotation mode (Dobin et al., 2013). Alignment, RNA-Seq, and Insert Size quality control metrics were generated using Picard v 2.10.10 (http://broadinstitute.github.io/picard). Homologous genes between vole, mouse, and human references were identified using biomart v2.56.1. When multiple mouse or human genes mapped to the same vole gene, the homologous gene with the highest percent identity to vole was selected.

##### Differential expression analysis

To remove non- and lowly-expressed genes, only genes with more than 2 counts per million (cpm) in at least 3 samples were retained. Principal component analysis of the cpm revealed that technical replicates all cluster tightly but showed a slight batch effect due to sequencing run (Fig. S5A). To remove this effect, we used RUVSeq v1.34.0 to estimate one factor of unwanted variation, W_1, which we later used to normalize the data prior to differential expression^125^. The factor analysis was performed on the deviance residuals from an initial generalized linear model regression of the upper quartile normalized counts on our covariates of interest (pair bonding status, genotype, sex). Expressed genes were tested for differential expression (DE), after combining technical replicates, with DESeq2 v1.40.2 using a design matrix that included the covariates and W_1^126^. The adaptive shrinkage estimator ashr v2.2-63 was used in order to be more robust to genes with low cpm values^127^. For DE analyses in general, while most changes were driven by female samples, the effect was maintained in the males at a smaller magnitude; thus, analyses were primarily performed for both sexes jointly to maximize accuracy and power.

###### Bonding effect

To compare pre- to post-pairing individuals, we performed the DE analysis in two ways. First, we compared pairing status regardless of genotype or sex to increase our statistical power and to identify a reproducible gene set of thirteen genes (Fig. 5B, 5C, S5B). While the relationship between the genes is unknown, there is support that the proteins are functionally related. Using the STRING protein database (STRINGdb_2.12.1), the human-homologs of the DE genes share more connections than we would expect by chance (Fig. S5C, p-value= 3.08E-0.7). Next, we compared pairing status using all three covariates, to identify subgroup-specific genes of interest and effect sizes (Fig. 5D, S5D). In both analyses, genes were considered significantly DE with a BH adjusted p-value < 0.05.

###### Genotype effect

To examine the effect of genotype, we first performed DE analyses between WT and Oxtr^−/-^ individuals, not including sex as a covariate to identify a reproducible and stringent gene set. We considered genes DE with an adjusted p-value < 0.05 and absolute log2 fold-change > 0.25 (Fig. 6B, S6A, S6B). To examine the functional relationship of the DE genes, we performed a Gene Ontology enrichment analysis using goseq v1.52.0 which controls for gene length bias^128^. We considered all expressed vole genes with a mouse homolog as our background and all DE genes with a mouse homolog as our foreground. GO categories with an adjusted p-value < 0.05 were considered significantly enriched. To help visualize the results, enriched GO categories were clustered according to their semantic similarity calculated with the R package GOSemSim v2.26.1 using method “Wang”^129^. Large, general, GO categories with over 1000 genes were excluded in visualizations (Fig. 6C). WikiPathways^130^ over-representation analysis was performed with R package clusterProfiler v4.8.3^131^ (Fig. S6D).

For disorder and disease set overrepresentation analysis, Chi-squared tests were used to assess whether Oxtr^1-/-^ disrupted genes occur in disease-associated gene lists at a frequency higher than we would expect by chance. Genes lists for autism spectrum disorder^74^, schizophrenia^75^, neurodevelopmental disorders^77^, and epilepsy^76^ were sourced and all alternate or outdated gene symbols were updated manually. For this test, we were interested in sex- and direction-specific enrichments, so we performed a DE analysis separately for males and females. Across all DE analyses, we defined DE genes more stringently (adjusted p-value < 0.01). For enrichment tests, our universe was all expressed vole genes with a human homolog. We split this analysis by fold-change direction to highlight the distinct enrichment in upregulated genes. P-values underwent BH correction across all 24 comparisons.

To compare to mouse neuronal populations, we leveraged single-cell data on mouse hypothalamus development^83^. The released Seurat object GSE132730_subset_cca_to_velo.rds.gz was accessed at GSE132730. Using the authors’ defined clusters (accessed at “prim_walktrap”), we assessed how similar each clusters’ gene expression is to the genes disrupted by the Oxtr^1-/-^ mutation. This was calculated as the Jaccard similarity coefficient using R package clustifyR v1.12.0^132^.

##### Weighted Gene Coexpression Network Analysis

###### Data

Starting with the RNAseq dataset of n=30 samples (one WT sample and 4 replicate samples were removed) and n=13357 genes, we removed genes with low expression (n=1738 genes with expression level in lowest quantile for all 30 samples) and/or low variability (n=8230 genes with coefficient of variation < 0.05), resulting in a trimmed dataset of n=3389 genes.

###### Identifying co-expression modules

We used the R package WGCNA (v1.70-3) to conduct WGCNA analyses^72^. Briefly, we calculated Pearson correlations for all gene pairs. Next, we constructed an unsigned weighted correlation network to identify co-expression modules comprised of highly correlated genes (positive or negative correlation) with high topological overlap^133^. Modules were defined as branches of a hierarchical cluster tree using the top-down dynamic tree cut method^72^. For each module, the expression pattern was summarized by the module eigengene (ME), which is defined as the right singular vector of the expression patterns. Pairs of modules with high module eigengene correlations (r > 0.75) were merged to maintain a level of decorrelation among MEs.

In more detail, a weighted unsigned network was computed based on a fit to scale-free topology, and the lowest thresholding power that resulted in a scale-free R^2^ fit of 0.9 was selected, and the pairwise topological overlap (TO) between genes was calculated^133^. We constructed a TO dendrogram using hierarchical clustering (R::hclust function, method= “average”), and defined modules using the WGCNA::cutreeHybrid function with minimum module size set to 30 genes. We then merged modules with high module eigengene correlations as describe above.

We calculated 3 sets of modules from different sample subsets:

1. Modules^WT^: n=15 modules constructed from n=18 WT samples with thresholding power β = 5.
2. Modules^Mut^: n= 43 modules constructed from n=12 Oxtr^1^-mutant samples with thresholding power β = 6.
3. Modules^All^: n=7 modules constructed from all n=30 samples with thresholding power β = 3

###### Relating modules to sample traits

In order to correlate modules with experimental conditions, we calculated the Pearson correlations between module eigengenes and sample metrics, including species (Mut=0, WT=1), condition (Pre=1, Post=0), sex (F=1, M=0), and quality control (QC) metrics (RIN, yield, sequencing run). We identified three modules that significantly correlated with a sample metric of interest but not with any of the QC metrics. Significant correlation was defined to be p_adj_ <= 0.05, where p_adj_ = BH correction by number of modules. These modules (M) were:

1. Mpink^WT^: positive correlation with pre-bonding status (Pearson R = 0.7, p_adj_ = 0.018)
2. Mbrown^All^: positive correlation with WT status (Pearson R = 0.55 , p_adj_ = 0.011)
3. Mturquoise^All^: positive correlation with WT status (Pearson R = 0.8, p_adj_ = 6.6e-7)

We calculated the module eigengenes (ME) for Mpink^WT^, Mbrown^All^, and Mturquoise^All^ for all n=30 samples using the WGCNA::moduleEigengenes function. We then conducted Wilcoxon rank sum tests (ggpubr::stat_compare_means function) to assess whether MEs were significantly different between the following groups:

1. MEpink^WT^: pre- versus post-bonded animals, separated by sex and species
2. Mbrown^All^: WT versus mutant animals, separated by sex and condition
3. Mturquoise^All^: WT versus mutant animals, separated by sex and condition

###### Comparing gene significance with module membership

We assessed whether genes whose expression highly correlate with module eigengenes of modules of interest (Mpink^WT^, Mbrown^All^, and Mturquoise^All^) are themselves highly correlated with traits of interest (bonding status or mutation status). Specifically, for the genes in Mpink^WT^, Mbrown^All^, and Mturquoise^All^, we calculated 1) gene significance (GS), defined as the absolute Pearson’s correlation between expression of a given gene and a trait of interest; and 2) module membership (MM), which is defined as the absolute Pearson’s correlation between the expression profile of a gene with the ME of a module to quantify the relationship between a gene and a given module^72^. We then calculated the Pearson correlations between:

1. GS^condition^ and MMpink^WT^ for Mpink^WT^ genes (R = 0.69, p = 5.5e-18)
2. GS^species^ and MMbrown^All^ for Mbrown^All^ genes (R = 0.63, p = 2.2e-32)
3. GS^species^ and MMturquoise^All^ for MMturquoise^All^ genes (R= 0.82, p<1e-200)

Hub genes were defined as those genes in the top 25% for both the absolute value of pink module membership and the absolute value of correlation with bonding status.

###### Assessing whether modules are enriched for DE genes

We were interested in assessing whether Mpink^WT^ genes are significantly enriched for any the DE pre vs post pairing gene set described above. We assessed for enrichment of DE gene sets using a hypergeometric test.

###### WT module preservation analysis (Zsummary)

We assessed how well modules^WT^ are preserved in mutant samples. We calculated a Z_summary_ statistic for each module^WT 134^ using the WGCNA::modulePreservation function with 200 permutations. The Z_summary_ measure combines module density and intramodular connectivity metrics to a composite statistic where Z>2 suggests moderate preservation and Z>10 suggests high preservation. We found of all the n=15 modules^WT^, Mpink^WT^ had the lowest Z_summary_ (Z_summary_=2) in mutant samples.

#### Histology

##### Immunohistochemistry (IHC)

Sexually naïve voles group housed by sex after weaning (P0, P21, adults:10 - 14 weeks old), were perfused with 25mL ice cold phosphate buffer (PBS) and fixed in 25mL of 4% paraformaldehyde in PBS (PFA). Whole brain was dissected and post fixed overnight at 4°C in 4% PFA in the dark. The tissue was cryoprotected for 24 hours in 30% sucrose in PBS and then embedded in OCT over dry ice to prevent cracking. 60μm serial sections from the entire PVN were collected in PBS and processed as free-floating sections. Sections were blocked using 10% donkey serum in 0.1% Triton X 100 in PBS and then stained with primary antibodies (Oxytocin Polyclonal Antibody, Invitrogen, Rabbit Polyclonal, Cat #PA5-26701, 1:1000 and Anti-Neurophysin 2/NP-AVP Antibody, clone PS 41, Millipore Sigma, Mouse Monoclonal, Cat#MABN845, 1:10,000) in 1% donkey serum with 0.1% Triton X 100 (staining buffer) in PBS at 4°C with shaking. The sections were rinsed and soaked in secondary antibodies (Cy™3 AffiniPure Donkey Anti-Rabbit IgG, Jackson ImmunoResearch Laboratory, Cat #711-165-152, 1:500 and Alexa Flour 488 Donkey Anti Mouse lgG, Thermo Fisher Scientific, Cat #A21202, 1:500) in staining buffer for 2 hours at room temperature. Sections were DAPI stained and transferred to glass slides in sequence and mounted in Aquamount. Slides were imaged using a confocal microscope (Zeiss LSM 700) and the zstacks were quantified (cell count by hand and area hand selected and measured) in ImageJ. Comparisons of cell density (count/area) were made in age and sex matched samples.

##### In situ hybridization (ISH)

ISH was performed using RNA probes prepared as described previously^135^, to detect expression of *Calcr, Dlk1*, and *Dgkb* mRNA in the adult vole brain. We prepared RNA sense and anti-sense probes corresponding to 706bp (CalcR), 773bp (Dgkb), and 796bp (Dlk1) for genes of interest identified in the WT vs Oxtr DEX list from our RNAseq data. For ISH on adult animals, 7-9 week old WT and Oxtr^1-/-^ voles of both sexes were either group housed with same sex conspecifics or paired for 4 days with a WT animal of the opposite sex following the timed mating protocol described above. The brains were processed as described for IHC, then embedded in OCT and sectioned at 50μm, mounted immediately after sectioning onto SuperFrost Plus glass microscope slides and stored at −80°C until ISH staining. Slides were fixed in 4% PFA for 20 minutes, rinsed in PBS, treated with proteinase K (10μg/mL, Roche) for 20 minutes, rinsed in PBS, and fixed again in 4% PFA for 5 minutes at room temperature. Slides were acylated for 10 mins then rinsed in 1% Triton100X in PBS followed by PBS. The slides were equilibrated in warm hybridization solution for 1-2 hours at 65°C and subsequently incubated with a temporary paraffin cover for 14 - 18 hours at 65°C in fresh hybridization buffer containing RNA probe. After incubation, the slides were dipped in 5X SSC warmed to 72°C to remove the parafilm coverslips and then washed in 0.2X SSC warmed to 72°C. The slides were blocked with 10% heat inactivated sheep serum (HISS) solution then stained for 12 - 18 hours at 4°C in buffer containing 10% HISS and alkaline phosphatase-conjugated sheep anti- digoxigenin antibody (1:2000, Roche). After extensive washing, the slides were incubated for ∼72 hours at 37°C in staining solution containing nitro blue tetrazolium and 5- bromo-4-chloro-3-indolyl-phosphate (Roche). The slides were finally washed, fixed in 4% PFA, and coverslipped. Slides were imaged using a Keyence fluorescence microscope and the z-stacks were quantified (cell count and area) in ImageJ. For signal quantitation, positive puncta were counted in every third coronal section encompassing the entire anterior to posterior gene expression domain. Cell counts were normalized by total area for a gene expression domain in each section. Positive signal was quantified by calculating total area of signal multiplied by the average intensity of signal (total area x mean pixel intensity) in gene expression domains of a section; every third section was quantified and added. Comparisons of cell counts were made in age and sex matched samples.

#### Quantification and Behavioral Analysis

For detailed statistical methods please see Table S1. The number of animals used was based on previous studies in the field by our group and others, combined with a power analysis. Assumptions of independence and normality were considered with multiple tests. Preference index (PI) was calculated as (partner huddle duration – stranger huddle duration)/(partner huddle duration + stranger huddle duration). Sliding partner preference was calculated by calculating PI for a single animal in 20-minute intervals where each interval overlapped the next by 15 minutes (e.g. 0-20mins, 5-25mins…). To look at convergence in sliding PI the area under the curve (Area above 0 – Area below 0, AUC) for each hour of the assay was determined for each animal and then pooled for each choosing animal by sex and then condition. Preference was determined as significantly convergent if the AUC from the hour of assay for all animals within the group significantly differed from 0 using a One sample t Test. The level of statistical significance for each test was set at p =< 0.05. Outliers were detected using a combination of z-score (+/− 3) and plots of original and log-transformed data. Analyses were completed in RStudio (version 2024.04.2+764) and GraphPad Prism (version 10.3.1)

