## Supplementary Figures for "Oxytocin receptor controls promiscuity and development in prairie voles"

### SUPPLEMENTAL DATA

**Supplementary Figure 1: Oxtr reduces the amount of time naïve animals take to show partner preference**

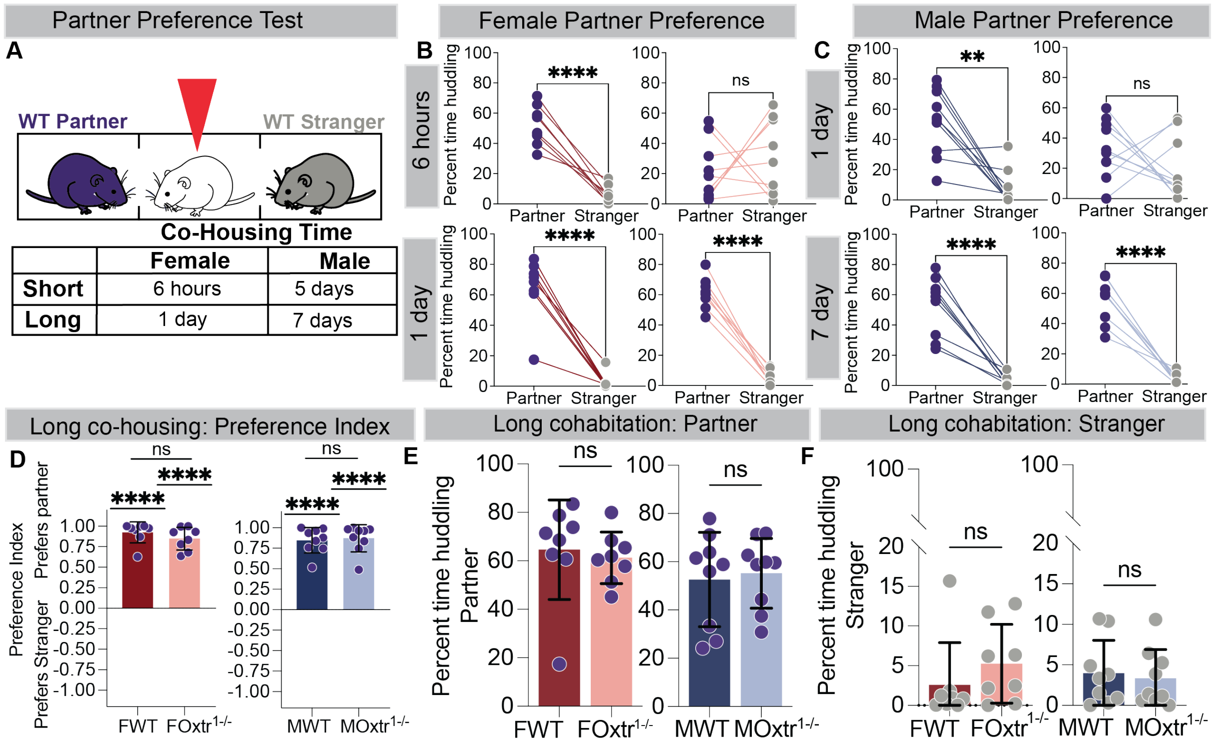


**Supplementary Fig 1: Oxtr reduces the amount of time naive animals take to show partner preference**

A. Schematic depiction of the PPT with the red animal indicating the subject animal. The assay was carried out on 2 independent cohorts at 2 time points: short and long, depending on the amount of cohabitation preceded the PPT.
B. Female Partner preference: Percent time spent with the partner was significantly greater than stranger except for $FOxtr^{1-/-}$ after short cohabitation.
C. Male Partner preference: Percent time spent with the partner was significantly greater than stranger except for $MOxtr^{1-/-}$ after short cohabitation.
D. A preference for partner is induced in all genotypes after a long cohabitation.

E. No difference in percent time spent huddling with partner between WT and $Oxtr^{1-/-}$ animals of either sex.

F. No difference in percent time spent huddling with stranger between WT and $Oxtr^{1-/-}$ animals of either sex.

See Table 1.

Mean +/- SD, n >8, *p =< 0.05, FWT = female WT, $FOxtr^{1-/-}$ = female $Oxtr^{1-/-}$, MWT = male WT, $MOxtr^{1-/-}$ = male $Oxtr^{1-/-}$.

**Supplementary Figure 2: Oxtr suppresses promiscuous behaviors in a state dependent manner in females**

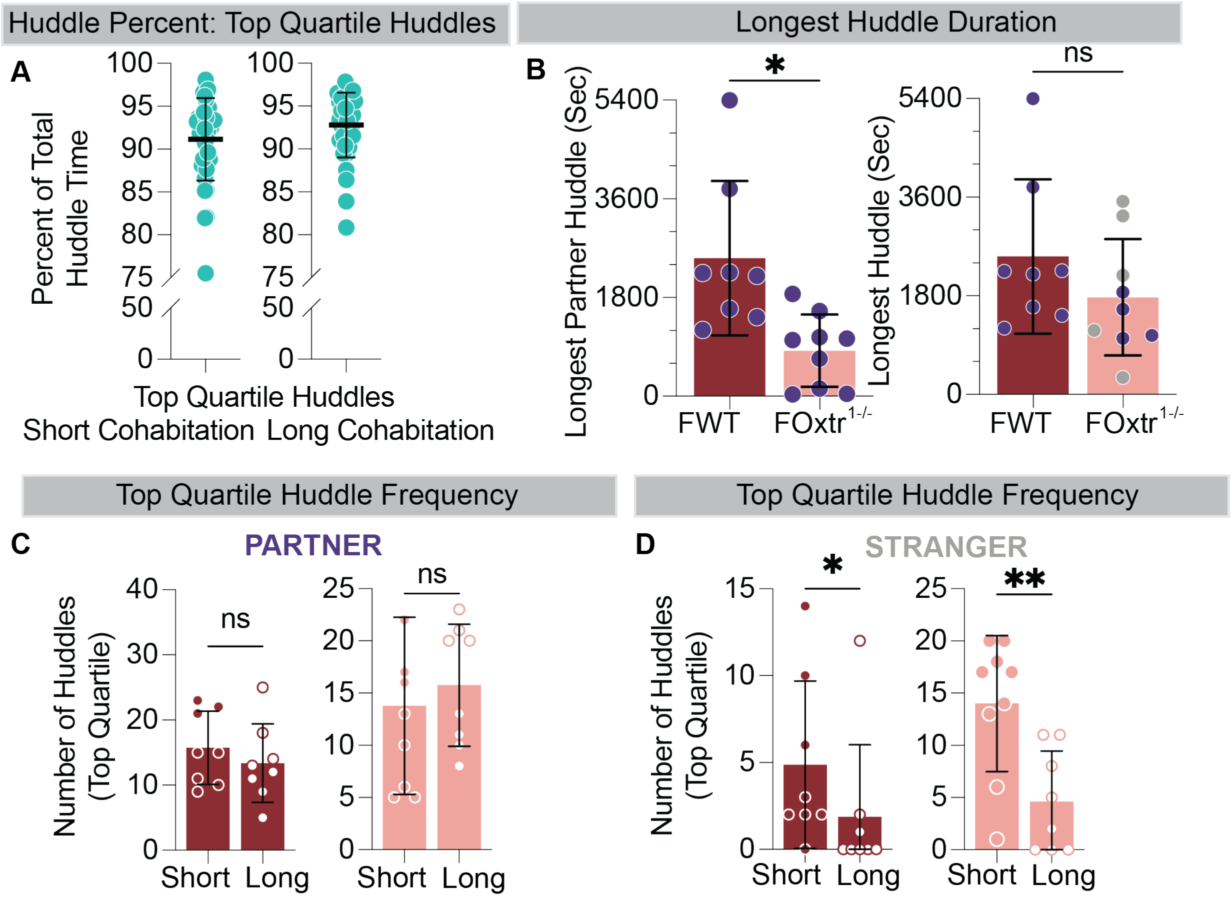


**Supplementary Fig 2: Oxtr suppresses promiscuous behaviors in a state dependent manner in females**

A. Each dot depicts the sum of the top quartile of huddles by duration as a percent of the total time spent in huddling with both partner and stranger. The average of all the individual top quartile huddle durations is greater than 90%. (Left: Short Co-housing, Right: Long Co-housing)

B. The longest huddle conducted by WT females with their partner is significantly longer than Oxtr1-/- females (left). There is no significant difference in the longest huddle overall (right).
C. Increasing cohabitation time does not change the frequency with which WT sibling controls nor $Oxtr^{1-/-}$ females huddle with partner.
D. Increasing cohabitation time decreases the frequency with which WT sibling controls and $Oxtr^{1-/-}$ females huddle with stranger.

See Table 1.

Mean +/- SD, *p =< 0.05, FWT = female WT, $FOxtr^{1-/-}$ = female $Oxtr^{1-/-}$, MWT = male WT, $MOxtr^{1-/-}$ = male $Oxtr^{1-/-}$

**Supplementary Figure 3: Oxtr function influences different prosocial behaviors across pair bonding in males**

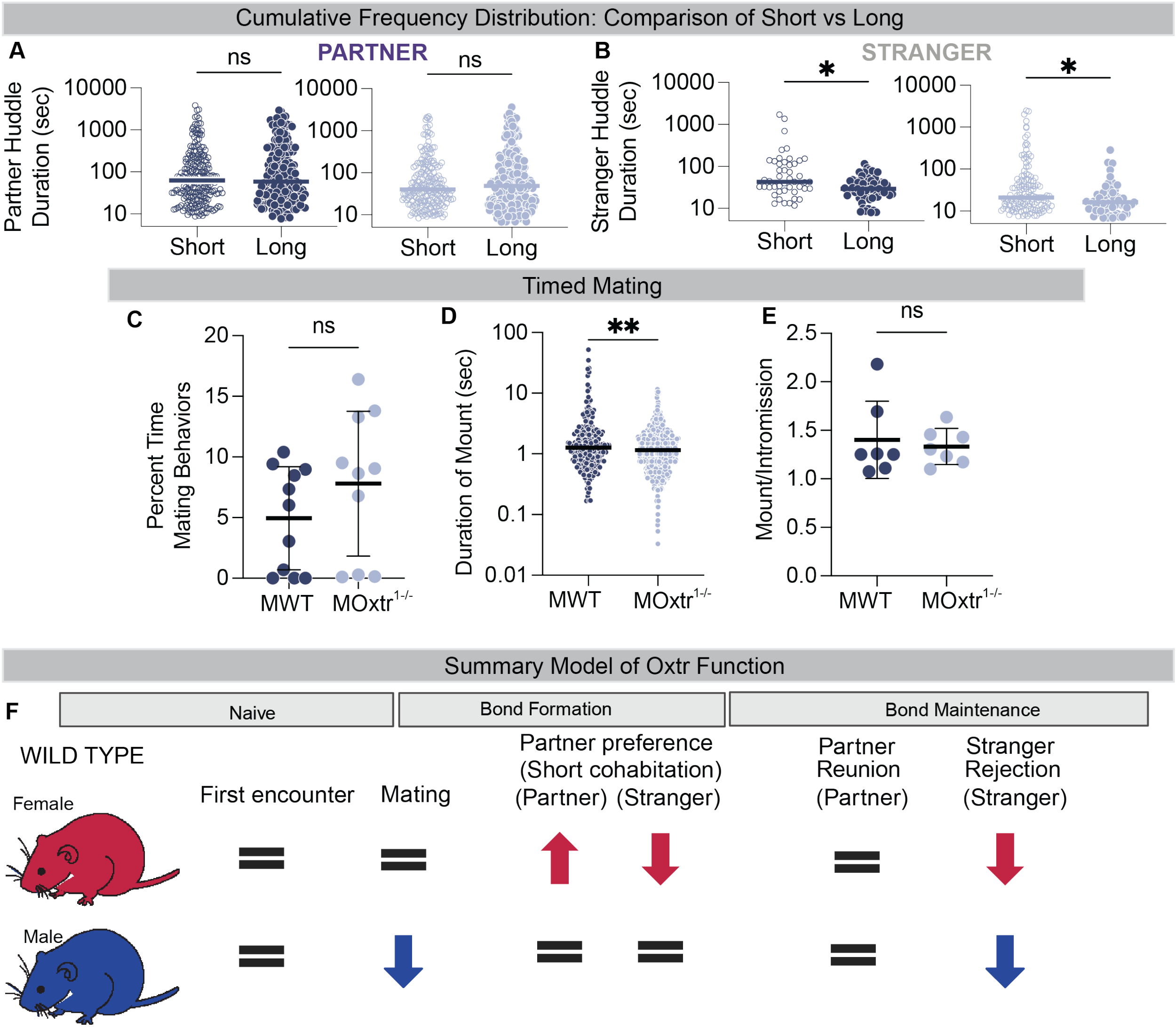


### Supplementary Fig 3: Oxtr function influences different prosocial behaviors across pair bonding in males

A. There is no significant change in the median huddle duration conducted by males with partners when cohabitation time is increased.

B. There is a significant decrease in the median huddle duration with stranger conducted by both WT and $Oxtr^{1-/-}$ males when cohabitation time is increased.

C. WT and Oxtr$^{1-/-}$ males spend the same amount of time performing mating behaviors.

D. The median duration of an Oxtr$^{1-/-}$ male mounting is significantly shorter than the WT sibling control mount duration.

E. Mount to Intromission ratio (counts) is the same for MWT and MOxtr$^{1-/-}$ animals

F. Timeline of bonding behaviors and their disruption in Oxtr$^{1-/-}$ animals. Arrows indicate difference from WT behavior and colored arrows show sex specific effects.

See Table 1.

Mean +/- SD, n >8, *p =< 0.05, FWT = female WT, FOxtr$^{1-/-}$ = female Oxtr$^{1-/-}$, MWT = male WT, MOxtr$^{1-/-}$ = male Oxtr$^{1-/-}$.

**Supplementary Figure 4: Oxtr sex-specifically influences social interactions between potential mates**

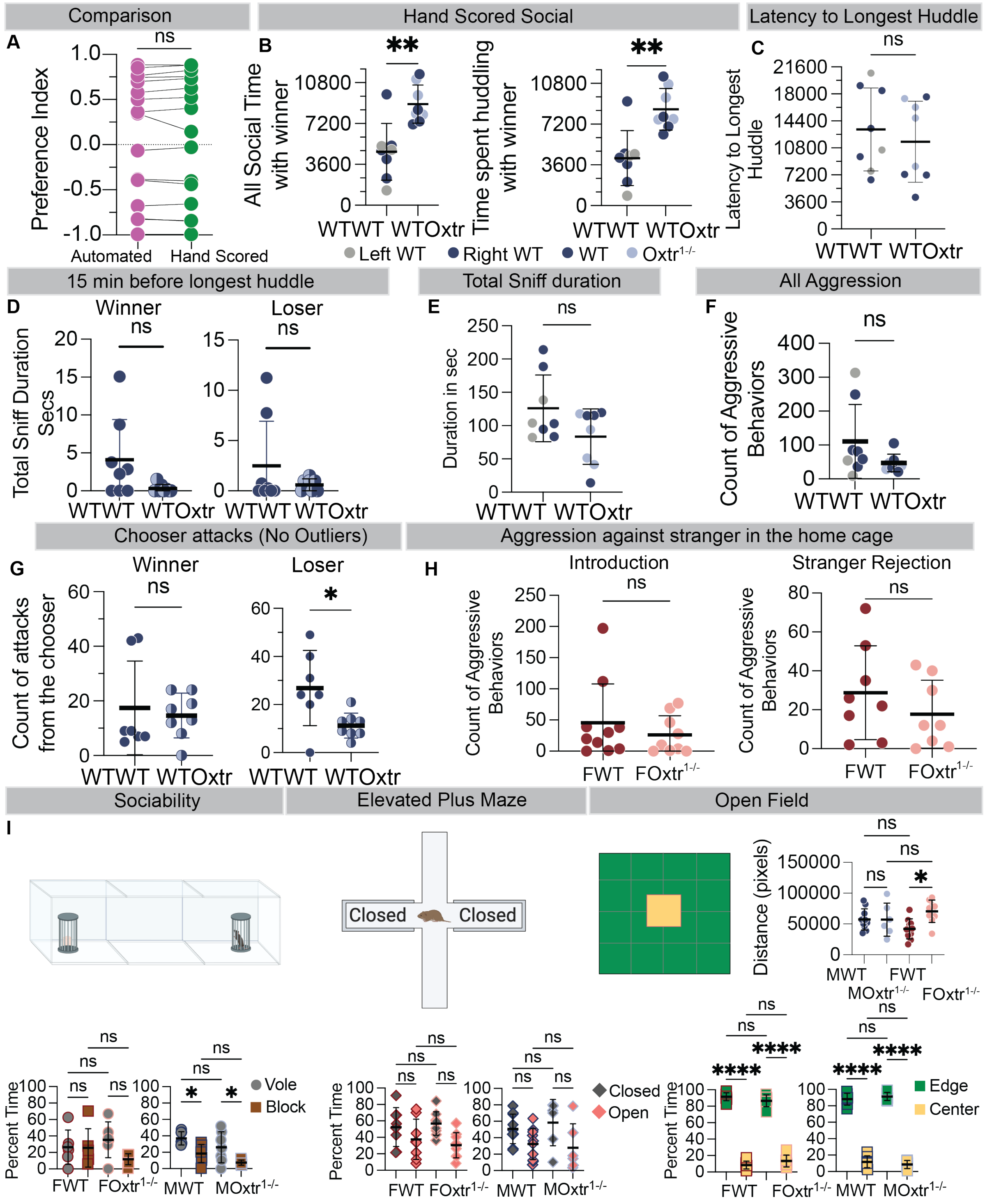


**Supplementary Fig 4: Oxtr sex-specifically influences social interactions between potential mates**

A. Comparison between automated scoring and hand scoring of the same footage shows no difference in the preference index.
B. WT female choosers spend more time performing social behaviors (left) and huddling (right) with the winner from WT-Oxtr lanes compared to the winner from WT-WT lanes. The color of the dot denotes the position (WT-WT) or the genotype (WT-Oxtr) of the winner.
C. There is no difference in the latency to start the longest huddle between WT-WT lanes and WT-Oxtr lanes. The color of the dot denotes the position (WT-WT) or the genotype (WT-Oxtr) of the winner.
D. There is no difference in the duration sniffing winners or losers by WT female choosers from WT-WT lanes and WT-Oxtr lanes 15 minutes before the onset of the longest huddle.
E. There is no difference in the total time spent by the WT female choosers sniffing the stimulus animals from WT-WT lanes and WT-Oxtr lanes.
F. There is no difference in the count of aggression (displayed + received) observed in WT-WT lanes compared to WT-Oxtr.
G. There is no difference in the number of attacks against the winner made by the WT female choosers from WT-WT lanes compared to WT-Oxtr lanes (left). WT female choosers from WT-WT lanes attack the loser more than WT female choosers from WT-Oxtr lanes (right).
H. There is no difference in the counts of aggressive behaviors (displayed + received) between WT females and $Oxtr^{1-/-}$ females against a WT stranger male during the initial introduction (left) nor post-bonding during stranger rejection.

I. $Oxtr^{1-/-}$ females move a greater distance than their WT siblings in the open field test. There is no difference between WTs and $Oxtr^{1-/-}$ in any other assay. (Sociability, elevated plus maze, open field)

See Table 1.

Mean +/- SD, n >6, *p =< 0.05, FWT = female WT, $FOxtr^{1-/-}$ = female $Oxtr^{1-/-}$, MWT = male WT, $MOxtr^{1-/-}$ = male $Oxtr^{1-/-}$.

### Supplementary Figure 5: Post pair bonding gene expression is driven by female specific changes dependent on Oxtr signaling

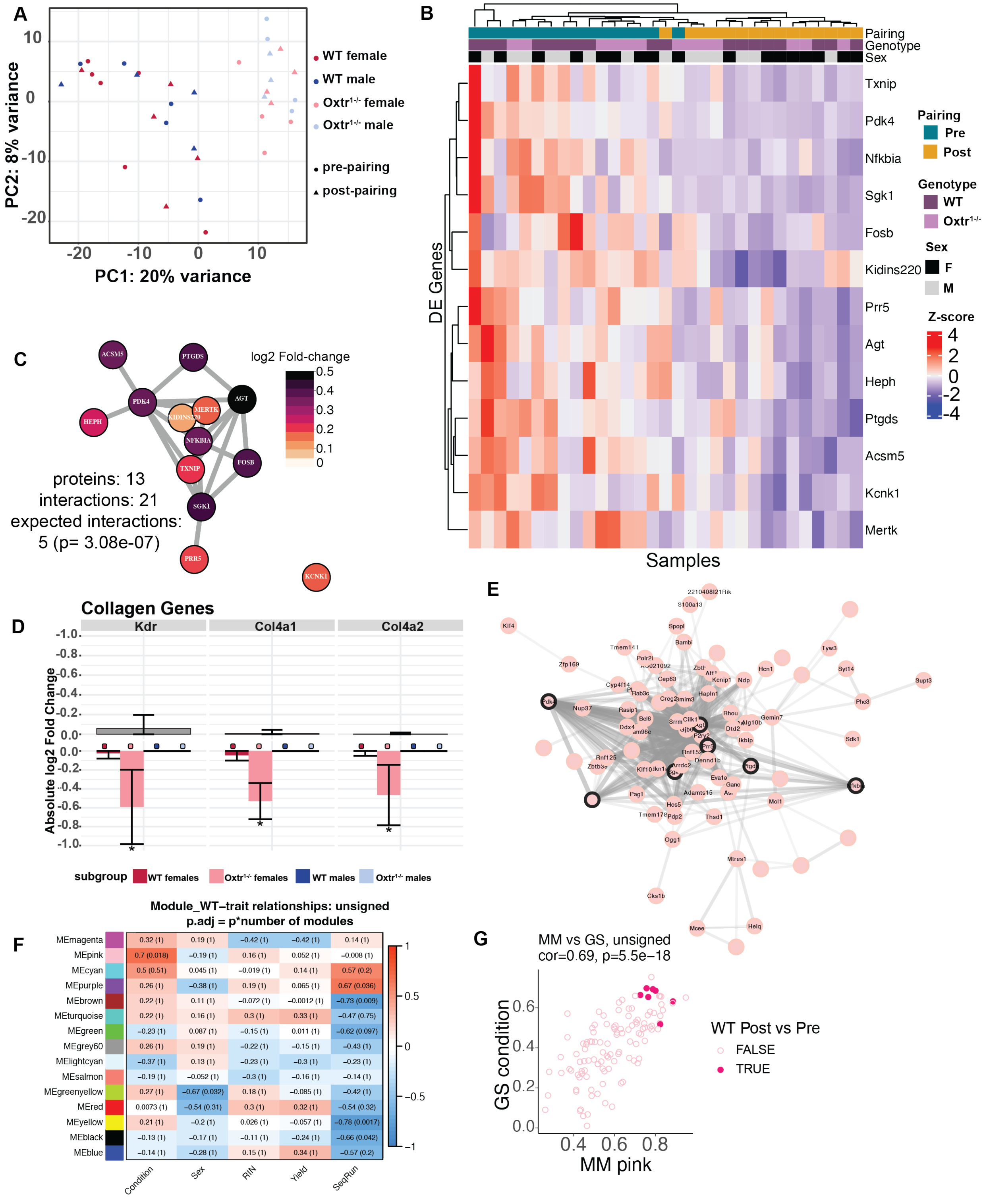

**Supplementary Fig 5: Post pair bonding gene expression is driven by female specific changes dependent on Oxtr signaling**

A. Principal component analysis (PCA) plot of the zero centered and normalized expression data shows separation by genotype. ($n$ = 19 pre-pairing and 15 post-pairing)
B. Gene expression heat map of DE genes between pre- and post-pairing conditions. DE genes (n=13) along the y-axis and samples (n=31) along the x-axis are ordered by hierarchical clustering. Cells are colored by the Z-score of normalized gene expression.
C. Protein-protein interaction network of DE genes across pairing conditions interact more than expected in STRING. Individual genes are shown as nodes and colored by $log_2$FC.
D. Bar graph structured like Figure 5D, but highlighting three collagen genes which are significantly downregulated specifically in post-pairing $Oxtr^{1-/-}$ females compared to pre-pairing.
E. $Mpink^{WT}$ interaction network consists of n=118 genes. Nodes are labelled by vole gene symbol (blank nodes do not have an associated gene symbol). Edge weights reflect topological overlap (TO), and only edges with TO>= 0.13 (~$95^{th}$ percentile of all edges) are shown. The n=7 of 13 WT_post_vs_pre DEX genes are outlined in black.
F. Heatmap showing the strength of Pearson correlations between MEs for $modules^{WT}$ sample metrics including species (Mut=0, WT=1), condition (Pre=1, Post=0), sex (F=1, M=0), and quality control metrics. (RIN, yield, sequencing run)

Heatmap color and first number reflect Pearson correlation coefficient, and the number in parentheses reflects adjusted p value. ($p_{adj}$ = p * number of modules)

G. The absolute values of $MMpink^{WT}$ and $GS^{Condition}$ for the n=118 genes in $Mpink^{WT}$ are significantly correlated (Pearson correlation = 0.69. p=5.5e-18).

**Supplementary Figure 6: Oxtr regulated genes are common to social behavior relevant processes across species and are similarly regulated across brain regions**

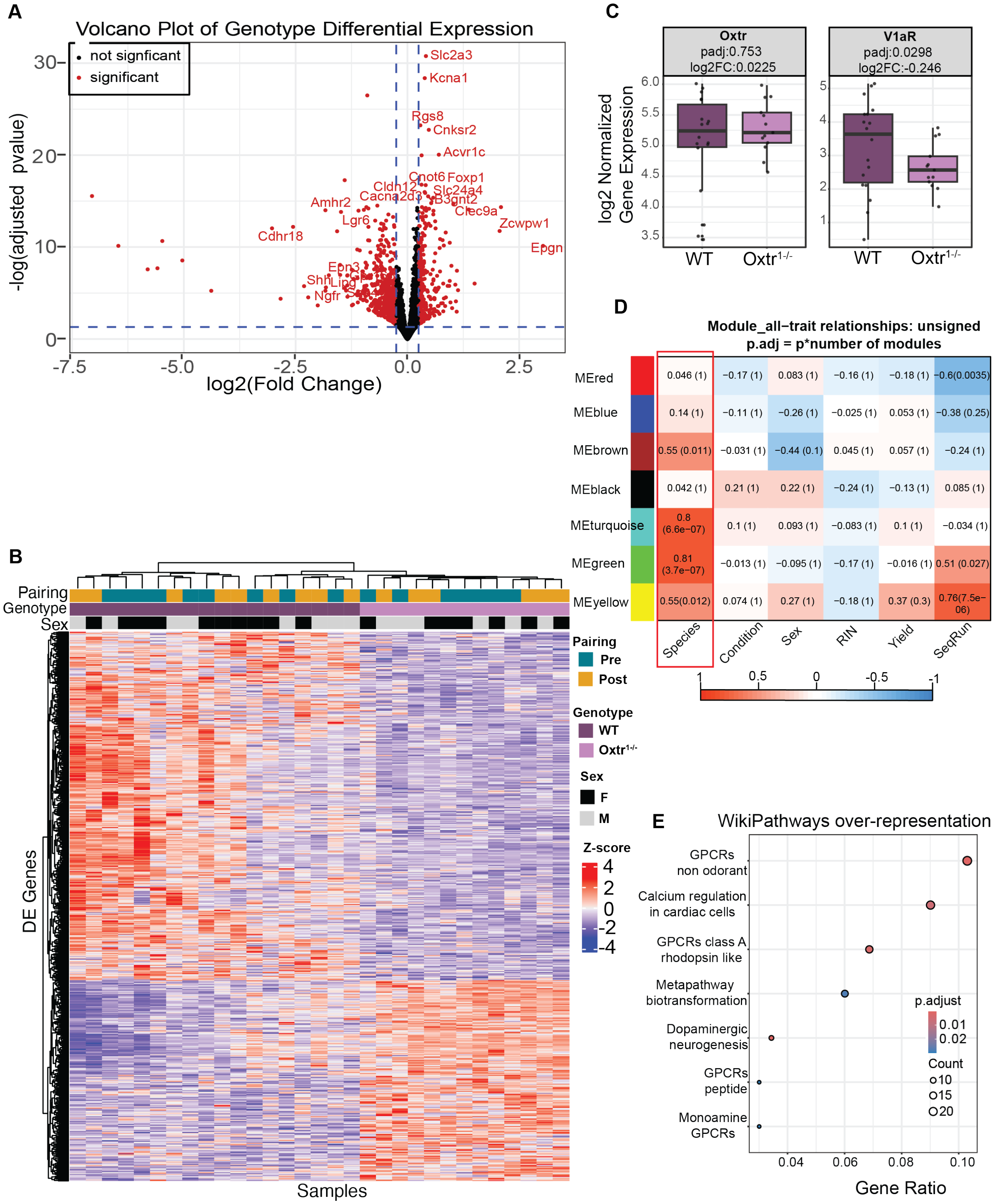

**Supplementary Fig 6: Oxtr regulated genes are common to social behavior relevant processes across species and are similarly regulated across brain regions**

A. Volcano plot of showing relationship of fold change to adjusted p-value when comparing Oxtr $^{1-/-}$; individuals to WT. Dashed blue lines show DE gene criteria thresholds at $p_{adj} < 0.05$ and absolute $log_2FC > 0.25$. Top DE genes with gene symbols are labeled on plot. For reference, positive fold changes are upregulated in Oxtr $^{1-/-}$ individuals.

B. Box plots of Oxtr and V1aR normalized gene expression across genotypes. Text displays $p_{adj}$ and $log_2FC$ values from comparison considering all WT and Oxtr$^{1-/-}$ samples.

C. Gene expression heat map of DE genes between WT and Oxtr $^{1-/-}$ samples. DE genes (n=1014, $p_{adj}$ <0.05, abs($log_2FC$) > 0.25) along the y-axis and samples (n=31) along the x-axis are ordered by hierarchical clustering. Cells are colored by the Z-score of normalized gene expression.

D. Heatmaps showing strength of Pearson correlations between MEs for modules$^{All}$ and sample metrics of interest, including species (Mut=0, WT=1), condition (Pre=1, Post=0), sex (F=1, M=0), and quality control metrics (RIN, yield, sequencing run). Heatmap color and first number reflect Pearson correlation coefficient, and the number in the parenthesis reflects adjusted p.value ($p_{adj}$ = p * number of modules). Red box indicates co-relation with genotype.

E. Over-representation analysis of genotype DE genes in biological pathways annotated in WikiPathways highlight processes related to GPCRs and dopaminergic neurogenesis. The x-axis is the ratio of DE genes in each category to the total number of DE genes in WIkiPathway database. The y-axis is all significant pathways (BH $p_{adj} < 0.05$). Dots are colored by adjusted pvalue gradient and size corresponds to the number of DE genes in pathway.

**Supplementary Figure 7: Developmental timeline for the effect of the loss of Oxtr on the cell density of OT and AVP positive cells in the PVN.**

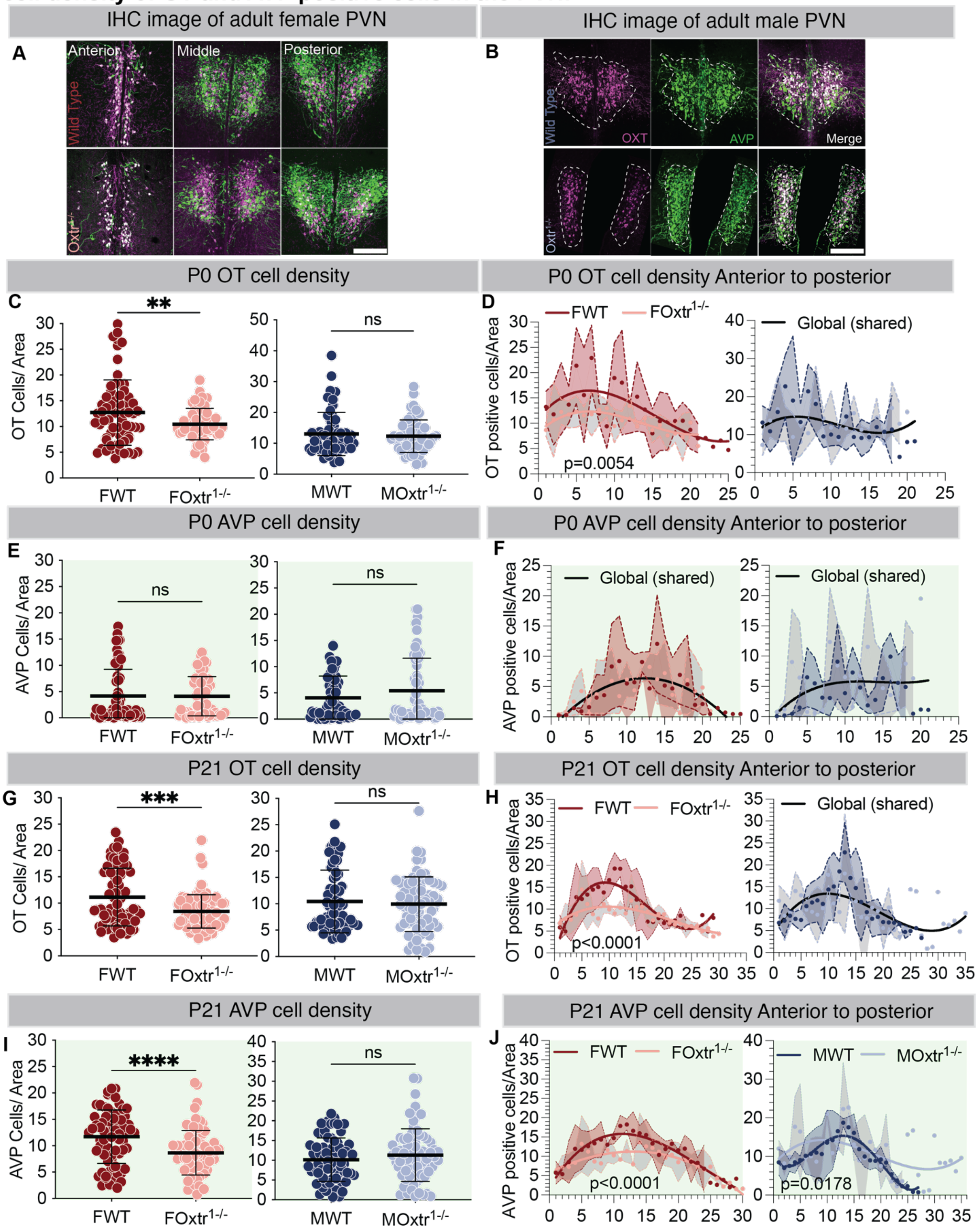

**Supplementary Fig 7: Developmental timeline for the effect of the loss of Oxtr on the cell density of OT and AVP positive cells in the PVN.**

A. Representative images from anterior, middle and posterior portions from adult WT and $Oxtr^{1-/-}$ female PVNs showing that cell density and PVN structure change based on the position within the PVN and genotype. The loss of Oxtr disrupts the structure of the PVN in the anterior regions more than the posterior.

B. Representative images from a WT and $Oxtr^{1-/-}$ adult male PVN showing the loss of cell density and disrupted structure in the anterior portion of the PVN.

C. PVN sections from P0 WT females have greater OT cell densities compared to sections from P0 $Oxtr^{1-/-}$ females (left). There is no difference in cell densities in PVN sections taken from P0 WT and $Oxtr^{1-/-}$ males (right).

D. Sections arranged from anterior PVN to posterior PVN show that the high OT cell density sections are lost in P0 $Oxtr^{1-/-}$ females, but not males.

E. – F. Similar analysis of P0 sections for AVP cell density shows no difference between WTs and $Oxtr^{1-/-}$s for either sex.

G. OT cell density comparison between WT and $Oxtr^{1-/-}$ sections taken from P21 animals. WT females have more high-density sections compared to $Oxtr^{1-/-}$ females.

H. Anterior to posterior classification of P21 OT sections from the PVN shows that $Oxtr^{1-/-}$ females, but not males, loose high-density sections from the anterior portion of the PVN.

I. – J. Similar analysis of AVP positive neurons from P21 animals shows the same result; a reduction in the high-density sections from the anterior portion of the PVN in Oxtr$^{1-/-}$ females. For AVP, there is also a structural difference in cell density between WT and Oxtr$^{1-/-}$ males.

See Table 1.

Mean +/- SD, n =3, *p =< 0.05, FWT = female WT, FOxtr$^{1-/-}$ = female Oxtr$^{1-/-}$, MWT = male WT, MOxtr$^{1-/-}$ = male Oxtr$^{1-/-}$, Scale bar = 100μM
